# Set mediates chromosome alignment and Cohesin cleavage independently of direct PP2A-B56-binding in oocyte meiosis

**DOI:** 10.64898/2026.08.20.745968

**Authors:** Leonor Keating, Arianna Esposito Verza, Warif El Yakoubi, Yulia Gryaznova, Safia El Jailani, Damien Cladière, Sandra Touati, Eulalie Buffin, Christophe Rachez, Varvara Sarli, Alberto M. Pendás, Wei Gu, Andrea Musacchio, Katja Wassmann

**Author notes:** Cell and Developmental Biology Center, National Heart, Lung, and Blood Institute, National Institutes of Health, Bethesda, MD 20894, USA.

## Abstract

Set promotes cohesion removal in mitosis by evicting phosphorylated Histone H1 and counteracting Sgo1. In addition, Set promotes chromosome alignment by counteracting Aurora B activation. The underlying molecular mechanisms through which Set performs these activities remain insufficiently characterized, but roles of Set as a Histone chaperone and PP2A inhibitor have been proposed. Building on our previous observations that Set promotes pericentromeric Cohesin removal in oocyte meiosis II, we generated an oocyte-specific conditional knock-out of Set to address its functions in meiosis. Similar to mitosis, Set depletion caused chromosome alignment and cohesion defects. We found that Set is required for accurate error correction by localizing Aurora B/C, and for efficient cleavage of the meiosis-specific Cohesin subunit Rec8 by Separase. Paired chromosomes and sister chromatids were often incompletely separated, likely a primary cause of missegregation. Set performed both its roles in a Sgo2-dependent manner, but, unexpectedly, independently of interaction with PP2A-B56. In line with a role of Set as a Histone chaperone, accumulation of phosphorylated Histone H1 in Set knock-out oocytes occurs concomitantly with reduction of oocyte-specific H1foo on chromosome arms, indicating that Set is required to create the optimal chromatin environment for efficient Rec8 cleavage by Separase in meiosis.

## Introduction

The timely removal of Cohesin, which holds sister chromatids together, is essential for generating two daughter cells with the correct genome content in both mitosis and meiosis. During mitosis, the majority of Cohesin located on chromosome arms is removed in prophase *via* the so-called prophase pathway, which involves the Cohesin-releasing activity of Wapl. Pericentromeric Cohesin, however, remains protected until all chromosomes are properly attached to the bipolar spindle and aligned at the metaphase plate. This ensures satisfaction of the spindle assembly checkpoint and activation of the protease Separase. Separase then cleaves the kleisin subunit Scc1 of the remaining pericentromeric Cohesin complexes, allowing the segregation of sister chromatids to opposite spindle poles^1–3^.

Mammalian oocytes undergo meiotic divisions after a prolonged dormant state in prophase I, known as the germinal vesicle (GV) stage. Upon hormonal stimulation, oocytes resume the first meiotic division, characterized by chromosome condensation and nuclear envelope breakdown (germinal vesicle breakdown, or GVBD). They then progress through meiosis I into meiosis II, arresting in metaphase II until fertilization triggers exit from meiosis II (**Figure 1A**). Prophase pathway-dependent removal of Cohesin does not play a role once oocytes resume meiosis I (e.g., after GVBD, which corresponds to release from prophase I arrest), even though complete loss of Wapl in mouse oocytes has been shown to increase Scc1-containing Cohesin on chromosome arms. Some precocious sister chromatid segregation in meiosis I has been observed in the absence of Wapl, which contradicts the idea that a Wapl-dependent prophase pathway is required for Rec8 Cohesin removal once oocytes enter the first meiotic division^4^.

**Fig. 1.**
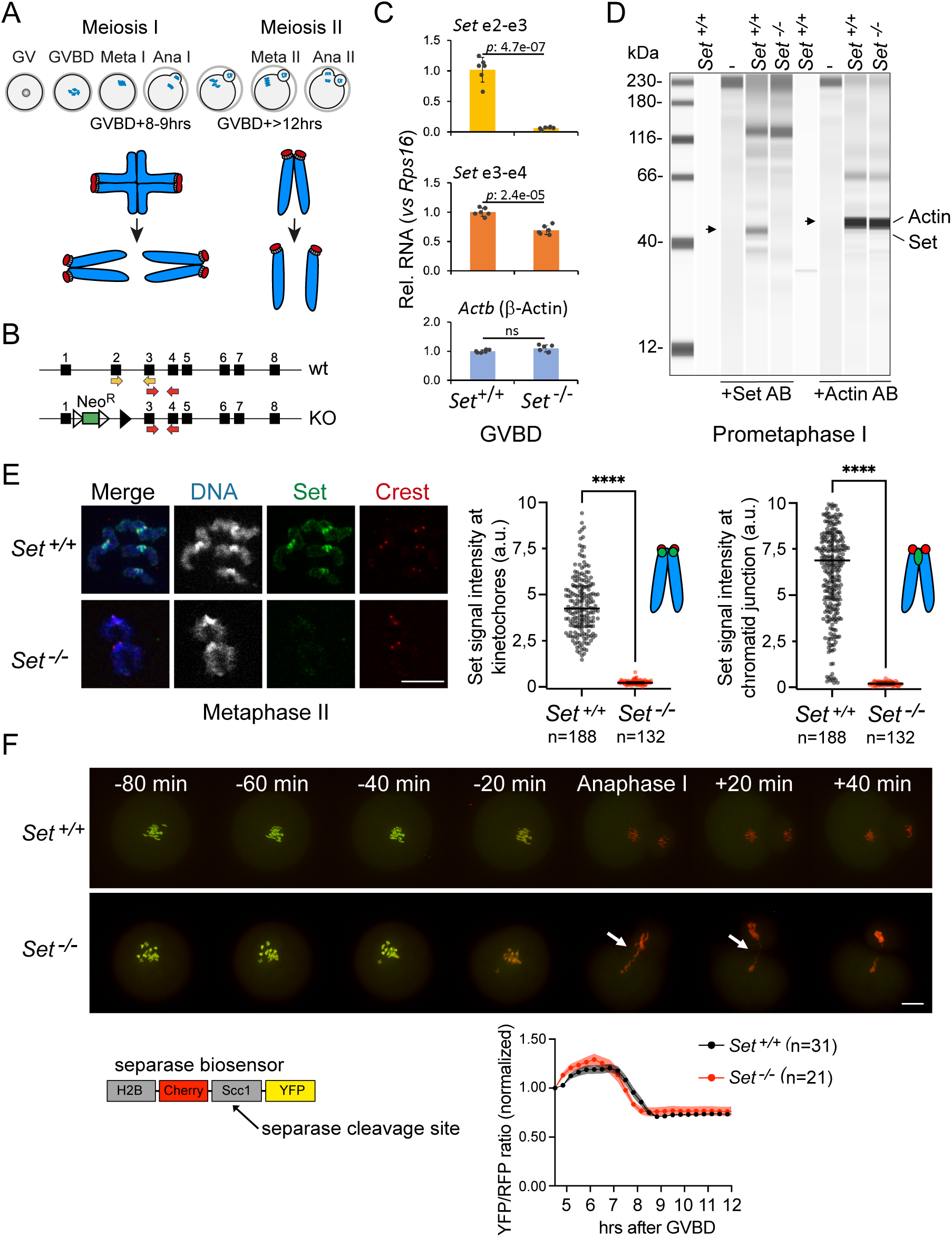
) Oocyte-specific conditional knock-out of *Set*. **A** Scheme of progression through meiosis I and II in mouse oocytes, and corresponding chromosome segregation patterns below. GV: Germinal vesicle state, corresponds to prophase I. GVBD: Germinal Vesicle Breakdown, marks resumption of meiosis I. Meta: metaphase, Ana: anaphase. Approximal times of metaphase I and metaphase II after GVBD are indicated. Of note, anaphase II occurs only upon fertilization or artificial override of CSF (Cytostatic Factor) arrest *in vitro*. **B** Knock-out strategy to invalidate Set in oocytes specifically, Exon 2 of *Set* is deleted in the knock-out allele. Primers used for RT-qPCR in (B) are indicated with yellow and red arrows. **C** RT-qPCR using *Set^+/+^* and *Set^-/-^* oocytes in GV stage, with primer pairs covering exon 2-3 and exon 3-4, indicated in yellow and red, respectively, in (A). β-Actin and Rsp16 were used as internal controls. **D** Visualization of protein levels present in 15 (to detect Actin) and 30 (to detect Set) prometaphase I oocytes (GVBD+ 4hrs) of the indicated genotype, using Jess™ capillary western blotting. Lanes without extract (-) and without antibody (indicated below) were used as controls. Actin was used as an internal control. **E** Metaphase II chromosome spreads from Set^+/+^ and *Set^-/-^* oocytes stained with anti-Set antibody (green), anti-Crest antibody to reveal kinetochores/centromeres (red) and Hoechst to stain DNA (blue). On the left, quantifications of Set signal normalized to Crest and the chromatid junction (see schemes, colours correspond to immunostaining). **F** Representative montage of selected time frames of live imaging movies of *Set^+/+^* and *Set^-/-^* oocytes progressing through meiosis I. Oocytes were injected with mRNA coding for a fluorescent Separase activity sensor, carrying a Separase cleavage fragment sandwiched between two fluorescent tags (YFP and Cherry) and localized to chromosomes due to Histone H2B (see scheme below). Separase activation is visualized due to change of colour of the sensor (from green to red). Arrow head indicates aberrant anaphase I in *Set^-/-^* oocyte. Below right: quantification of YFP/RFP ratio as a read-out of Separase activity towards the sensor. Scale bars: (F): 20 μm, (E): 10 μm. In (C), dots indicate individual experimental repeats, and unpaired student t-test was used. For quantifications in (D), n indicates the number of dyads analyzed, median and interquartile range are indicated and **** corresponds to p < 0,0001, using Mann-Whitney U test. In (E), each dot is mean and shaded areas indicate +/- SD. At least three independent biological repeats were analyzed for each condition.

Crucially, correct chromosome and sister chromatid segregation in meiosis I and II, respectively, require the stepwise removal of cohesion-from chromosome arms in meiosis I and from the pericentromere in meiosis II. This stepwise removal is essential for the segregation of paired chromosomes in meiosis I and paired sister chromatids in meiosis II^5^. Although Cohesin with distinct kleisin subunits (Rec8 and Rad21L) coexists in mouse oocytes, only Rec8-containing Cohesin is essential for cohesion^6^. Rec8-Cohesin is removed through cleavage by Separase in meiosis I and II^6–9^. In meiosis I, Separase can cleave phosphorylated Rec8 on chromosome arms only, as Rec8 at the pericentromere is protected through the recruitment of Sgo2, which localizes PP2A-B56 to the centromere region. There, PP2A-B56 is thought to counteract phosphorylation of Rec8 by Aurora kinases, which is necessary for efficient cleavage^10^. Intriguingly, PP2A and Sgo2 are prominently localized to the centromere region in oocyte meiosis II^11–13^, though their role there remains unknown and may be related to other functions, such as promoting proper attachments and alignment^14^. Our recent data show that pericentromeric Cohesin protection is absent in meiosis II^15^, but how PP2A and Sgo2 are prevented from protecting pericentromeric Cohesin in metaphase II remains unclear.

The histone chaperone Set has been shown to be required for chromosome alignment and proper prophase pathway-dependent and Separase-dependent removal of Cohesin and thus, chromosome segregation, in mitosis. Set is thought to contribute to the removal of histone H1 phosphorylated on serine/threonine 18 (depending on the isoform, H1p18) and Sgo1, thereby allowing Cohesin removal by the prophase pathway and Separase^16–18^, but the underlying molecular mechanisms are still unknown. Set has been attributed multiple functions, including acting as a histone chaperone, a PP2A inhibitor, a component of the INHAT (Inhibitor of Histone Acetyltransferases) complex, and a regulator of transcription^19, 20^. It is also required for correct acetylation of histone and non-histone proteins such as p53 and Foxo1^19, 20^. Set is thought to regulate the Aurora B-dependent error correction pathway in a Sgo2-and PP2A-dependent manner, thus promoting chromosome alignment and proper microtubule attachments^17, 21^.

Set was also identified as an interaction partner of Sgo2 in oocyte meiosis^12, 22^. Morpholino-mediated, transient knockdown of Set leads to defects in sister chromatid segregation in oocyte meiosis II, whereas overexpression of Set causes precocious sister chromatid segregation in meiosis I. Since protection of pericentromeric Cohesin in oocytes requires the recruitment of PP2A-B56 through Sgo2, we and others have proposed that Set may counteract Cohesin protection by directly inhibiting PP2A at the centromere^12, 23, 24^. However, no *in vivo* data have demonstrated that Set can indeed interfere with PP2A-B56 substrate dephosphorylation at the pericentromere to remove protection, leaving open the possibility that Set promotes sister chromatid segregation through a different mechanism.

Here, we employed a conditional, oocyte-specific knockout approach to clarify Set’s role in Cohesin removal and chromosome alignment during oocyte meiosis. Our study shows that oocytes lacking Set exhibit defects in proper chromosome alignment in meiosis I due to failures in error correction. Additionally, Cohesin is not efficiently removed from chromosome arms in meiosis I, leading to the presence of chromosome pairs (bivalents) and sister chromatids with fused chromatid arms (“zipped dyads”) in meiosis II. We characterized functional domains in Set required for interaction with PP2A and Sgo2 *in vitro* and defined which domains are required for chromosome alignment and Cohesin removal through rescue experiments in oocytes. Our data show that both alignment and Cohesin removal are mediated by Set in a Sgo2-dependent manner, but independently of its interaction domain with PP2A-B56. Importantly, we found that Set contributes to the correct chromatin environment for optimal Rec8 cleavage by promoting incorporation of the oocyte-specific histone H1foo (also called H1oo) instead of somatic linker histones.

## Results

### Generation of an oocyte-specific Set knock-out mouse model

In humans and mice, there are two nearly identical isoforms of Set, designated α and β (also referred to as Set 2 and 1 or Set 201 and Set 202, respectively), which differ only in their very N-terminal 25 amino acid sequence in exon 1 (**Supplementary Figure S1A**). Both isoforms are transcribed and present in mouse oocytes (**Supplementary Figure S1B**). When overexpressed during oocyte meiosis I, both isoforms induce precocious sister chromatid segregation, with Setβ consistently producing a stronger phenotype (**Supplementary Figure S1C**), in line with published data^23^.

Given Set’s multiple roles in essential mitotic events, complete loss of Set is expected to affect overall survival in a knock-out mouse model. Thus, to study Set’s role specifically in female meiosis, we generated an oocyte-specific conditional knock-out using a strain with exon 2 of the *Set* allele flanked by *loxP* sites (*Setfl/fl*)^25^, crossed with a strain harboring Cre recombinase under the control of the oocyte-specific *Zona pellucida 3* (*Zp3*) promoter^26^. The resulting knock-out allele lacked exon 2 (**Figure 1B**). Importantly, both Set isoforms were targeted by this knock-out strategy.

Oocytes were retrieved from *Setfl/fl* and *Setfl/fl Cre+* females, induced to resume meiosis I, and allowed to undergo meiotic maturation. No full-length Set mRNA was detected in oocytes from *Setfl/fl Cre+* females (**Figure 1C**). Set protein (full-length or truncated) was undetectable by Jess capillary Western blot from whole oocyte extracts and on chromosome spreads using an antibody directed against the C-terminus of Set (**Figures 1D and 1E**)^27^. Thus, oocytes from *Setfl/fl Cre+* females lacked detectable full-length Set mRNA or Set protein (truncated or full-length) and were designated *Set^-/-^* oocytes.

### *Set^-/-^* oocytes progress through meiosis I without delay, but missegregate chromosomes

To determine whether oocytes lacking Set exhibit defects in meiotic cell cycle progression, we expressed a Separase biosensor by injecting control and *Set^-/-^* GV-stage oocytes with mRNA encoding the sensor. The sensor contains a Separase cleavage site and two fluorescent tags, whose colocalization is lost upon Separase activation and subsequent cleavage (scheme in **Figure 1F**). Chromosomal localization is ensured by a Histone H2B tag, which remains associated with chromosomes even after cleavage. This allows simultaneous tracking of chromosome movements and Separase activity *via* live imaging^28^. We found that Separase was activated on time in both control and *Set^-/-^*oocytes. However, chromosomes appeared poorly aligned, and missegregation events were frequently observed in *Set^-/-^* oocytes (**Figure 1F**). The absence of delay in Separase activation indicated that the spindle assembly checkpoint was not activated in oocytes lacking Set, despite the misaligned chromosomes.

### Kinetochore attachment errors in *Set^-/-^* oocytes are not corrected

The misaligned chromosomes in *Set^-/-^* oocytes led us to investigate whether kinetochores were properly attached to stable microtubule fibers. Cold treatment of oocytes prior to fixation allows staining of stable microtubules that are end-on attached to kinetochores. Whole-mount immunofluorescence of *Set^-/-^* oocytes revealed that kinetochores were attached; however, chromosomes were often found along the spindle and even at spindle poles. This phenotype was attributable to Set loss, as injecting GV oocytes with mRNA encoding GFP-Setβ largely rescued the chromosome alignment defects (**Figure 2A**, see also below, **Figure 3E**).

**Fig. 2.**
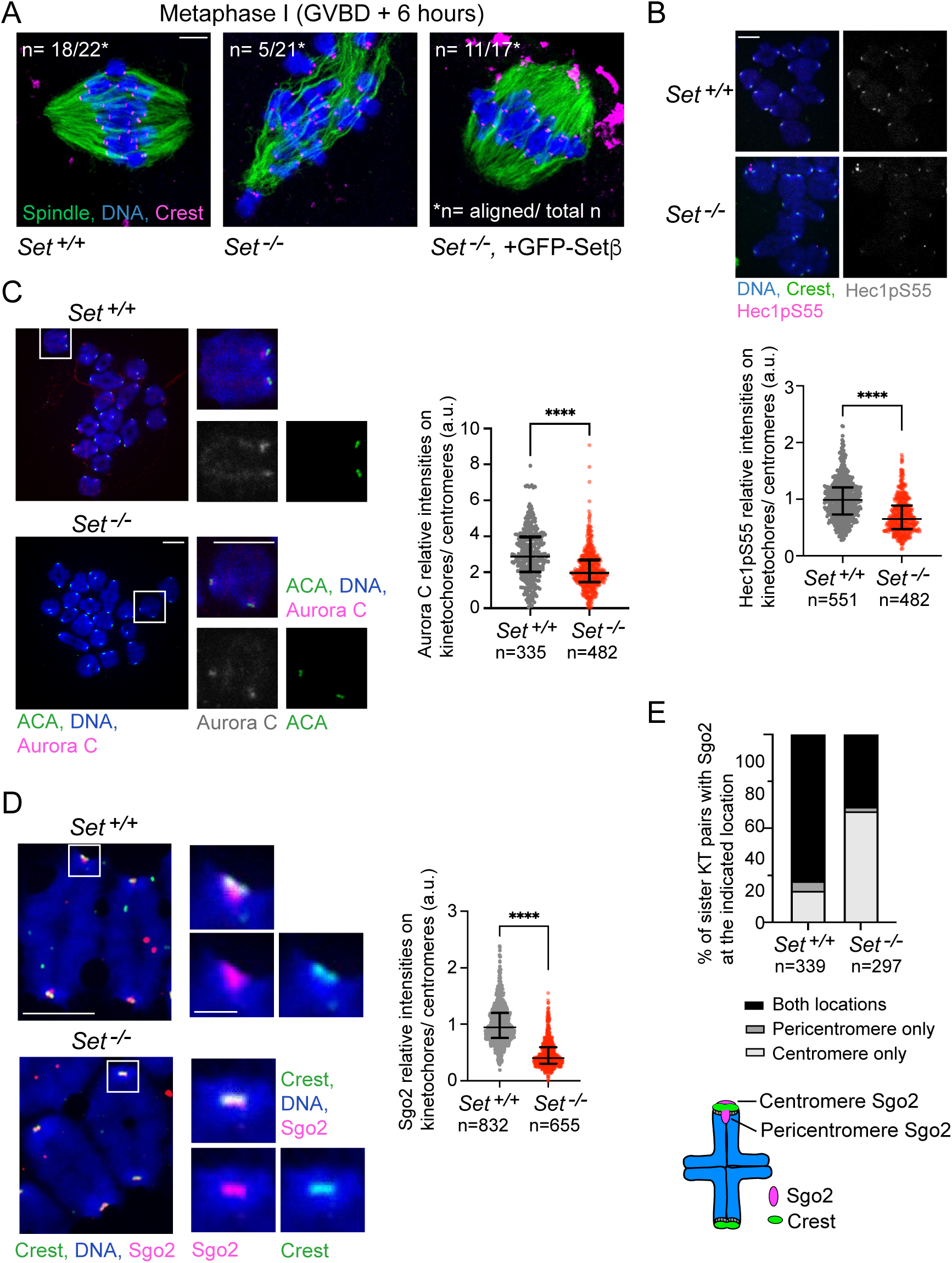
) Error correction pathway deficient in oocytes without Set. **A** Whole mount immunofluorescence staining of *Set^+/+^* and *Set^-/-^* oocytes in metaphase I, injected with mRNA encoding wild type Setβ where indicated. Cold-treated oocytes were stained with anti-Crest (red, kinetochores) and anti-α-tubulin antibody (green, spindles), as well as Hoechst (blue, chromosomes). The number of oocytes with aligned chromosomes and corresponding to the representative images shown from the total number of oocytes is indicated as n/total n. **B** Chromosome spreads of *Set^+/+^* and *Set^-/-^* oocytes in metaphase I, stained with anti-Hec1pS55 antibody (red), anti-Crest antibody to reveal kinetochores/centromeres (green) and Hoechst to stain DNA (blue). Below, quantifications of the Hec1pS55 signal relative to Crest signal. n indicates number of kinetchore pairs analysed. **C** Chromosome spreads of *Set^+/+^* and *Set^-/-^* oocytes in metaphase I, stained with anti-Aurora C antibody (red), anti-ACA antibody to reveal kinetochores/centromeres (green) and Hoechst to stain DNA (blue). Zoom of selected kinetochore pairs is shown. On the right, quantifications of the Aurora C signal overlapping with ACA. n indicates number of kinetchore pairs analysed. **D** Chromosome spreads of *Set^+/+^* and *Set^-/-^* oocytes in metaphase I, stained with anti-Sgo2 antibody (red), anti-Crest antibody to reveal kinetochores/centromeres (green) and Hoechst to stain DNA (blue). Zoom of selected kinetochore pairs is shown. Red arrow indicates Sgo2 at the pericentromere. On the right, quantifications of the Sgo2 signal relative to Crest signal. n indicates number of kinetchore pairs analysed. **E** Percentage of kinetochore pairs with Sgo2 at the centromere and/or the pericentromere. See scheme on the left for classification. For quantifications in (B and C), n indicates the number of dyads analyzed, median and interquartile range are indicated and **** corresponds to p < 0,0001, using Mann-Whitney U test. At least three independent biological repeats were analyzed for each condition. Scale bars: (A): 5 μm, (B-D): 10 μm, insert in (D): 2 μm.

**Fig. 3.**
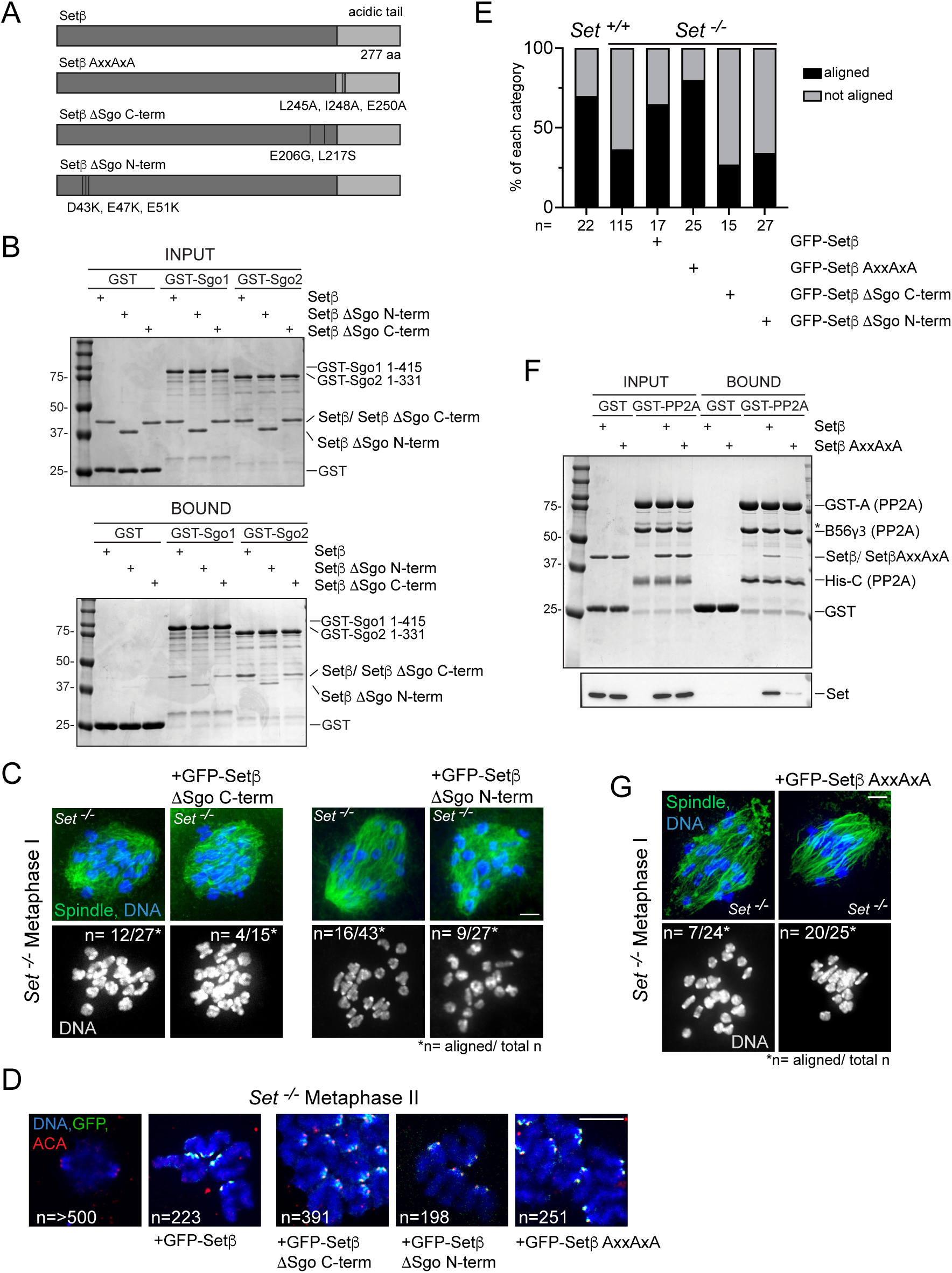
) Set promotes chromosome alignment independently of PP2A-B56 SliM, but requires interaction domains for Shugoshin. **A** Scheme of the Set mutants used. Amino acid positions correspond to mouse Setβ. **B** Binding assay on Glutathione beads with GST-Sgo1 1-415, GST-Sgo2 1-133 (bait) and GST (negative control). Setβ wild-type, Setβ ΔSgo C-term or Setβ ΔSgo N-term were added in solution such as indicated. Beads were recovered by centrifugation, washed and analyzed by SDS-PAGE and coomassie blue staining. The experiment consists of two biological replicates. **C** Whole mount immunofluorescence staining of *Set^+/+^* and *Set^-/-^* oocytes in metaphase I, injected with mRNA encoding for Setβ ΔSgo C-term or Setβ ΔSgo N-term such as indicated. Cold-treated oocytes were stained with anti-α-tubulin antibody (green, spindles), as well as DAPI (blue, chromosomes). The number of oocytes with aligned chromosomes and corresponding to the representative images shown from the total number of oocytes is indicated as n/total n. **D** GV-stage *Set^-/-^* oocytes were injected with mRNAs coding for the indicated GFP-tagged Setβ mutants, induced to resume meiosis I, and fixed for chromosome spreads in metaphase II. Spreads were stained with anti-GFP antibody to reveal exogenously expressed Set (green), anti-ACA antibody to visualize kinetochores/centromeres (red) and DAPI to stain DNA (blue). n indicates the number of chromosomes analyzed, from three independent biological repeats per condition. **E** Percentage of *Set^+/+^* and *Set^-/-^* oocytes of each category (chromosomes aligned, not aligned) from panels (C and E). n indicates number of oocytes analyzed for each condition from at least three independent biological repeats. **F** Binding assay on Glutathione beads with GST-PP2A-B56γ3 holoenzyme (bait) and GST (negative control). Setβ wild-type or Setβ AxxAxA were added in solution. Beads were recovered by centrifugation, washed and analyzed by SDS-PAGE (upper square, Commassie blue staining) or Western blotting against SET protein (lower square). The asterisk marks a contaminant of the GST-A subunit in the input fractions. The experiment consists of two biological replicates. **G** Whole mount immunofluorescence staining of *Set^+/+^* and *Set^-/-^* oocytes in metaphase I, injected with mRNA encoding for GFP-Setβ AxxAxA where indicated. Cold-treated oocytes were stained with anti-α-tubulin antibody (green, spindles), as well as DAPI (blue, chromosomes). The number of oocytes with aligned chromosomes and corresponding to the representative images shown from the total number of oocytes is indicated as n/total n. Scale bars: (C): 5 μm, (D): 10 μm, (G): 5 μm.

Misaligned chromosomes are typically recognized by the Aurora B-dependent error correction pathway due to imbalanced tension applied by the bipolar spindle on a given kinetochore pair. Aurora B and the oocyte-specific Aurora C phosphorylate Hec1 to destabilize erroneous attachments, allowing reformation of proper attachments^29–32^. However, in *Set^-/-^* oocytes, we observed reduced Hec1 phosphorylation at serine 55-a readout for ongoing error correction-appeared reduced compared to control oocytes (**Figure 2B**). This suggests that error correction is diminished or absent in *Set^-/-^* oocytes, explaining why chromosomes are misaligned. Similar to mitosis, where Set was proposed to inhibit PP2A to facilitate kinase activity of Aurora B^17^, this suggests that Set is required for a balanced activity of Aurora B/C also in meiosis.

### Reduction of Aurora C at centromeres in *Set^-/-^* oocytes

To understand why there was less ongoing error correction and misaligned chromosomes in *Set^-/-^* oocytes, we asked whether Set is required for proper localization of Aurora B/C to centromeres. Chromosome spreads in metaphase I were stained for Aurora C, and the signal overlapping with the centromere signal was quantified. Of note, Aurora C is also localized to the inter-chromatid axis^33, 34^. Crucially, Aurora C at centromeres was significantly reduced in *Set^-/-^* oocytes compared to controls (**Figure 2C**). Thus, error correction in absence of Set may be less efficient due to reduced recruitment of Aurora kinases.

### Reduction of a specific pool of Sgo2 in *Set^-/-^* oocytes

Sgo2 centromeric localization in mitosis and meiosis depends on Aurora B/C, Bub1, and Mps1 kinase activities^35–38^. There, Aurora B/C phosphorylates Sgo2, converting it to an inhibitor of its own kinase activity^35, 39^. Conversely, Sgo2 was also found to recruit Set to inner centromeres in mitosis, but this time, to promote Aurora B activity by inhibiting PP2A^17^. Sgo2 depletion in oocytes promotes Aurora C activity without affecting Aurora C localization^39^. Thus, it does not phenocopy the effects on Aurora kinase activity observed here upon depletion of Set-it rather results in opposite outcomes. To gain further insights we investigated whether Sgo2 localization is affected in oocytes without Set.

Indeed, metaphase I spreads revealed that Sgo2 levels were reduced in oocytes lacking Set (**Figure 2D**). Of note, there are at least two distinct pools of Sgo2 around centromeres in meiosis I: one at the pericentromere and one at the centromere. Counterintuitively, in the mouse, the centromeric pool (overlapping with CREST staining) is required for pericentromeric Cohesin protection in oocytes^11, 36^. It was foremost the pericentromeric pool - the one not required for Cohesin protection-that was lost in *Set^-/-^* oocytes (**Figure 2E**). Thus, Set is indeed contributing to Sgo2 localization at the pericentromere, probably through recruitment of centromeric Aurora C.

### Set interacts with Sgo2 through its N- and C-terminal domains for chromosome alignment

Two Set mutants impaired in the interaction with Shugoshins have been described. Mutations in the C-terminal region of Set (Setβ ΔSgo C-term) were proposed to affect mainly interaction with Sgo2^17^. Mutations in the N-terminus of Set (Setβ ΔSgo N-term) were proposed to impair interactions with both Sgo1 and Sgo2 (**Figure 3A**)^18^. Due to the limited protein yield from mouse oocytes (less than 25 ng per oocyte), we performed *in vitro* binding assays with purified components under high salt conditions to compare both mutants and their ability to bind Sgo1 and Sgo2. Both Setβ ΔSgo C-term and Setβ ΔSgo N-term proteins exhibited reduced binding to truncated GST-Sgo2 compared to wild-type Setβ. Wild-type Setβ bound Sgo2 more strongly than Sgo1, and the interaction was affected in a similar manner by mutating either interaction domain (**Figure 3B and Supplementary Figure S2A**). At physiological salt concentrations Setβ bound equally efficiently to Sgo1 and Sgo2. Also under these conditions, the Setβ ΔSgo mutants did not impair binding, suggesting that their binding affinity remains too high for assessing the functionality of these Sgo interaction domains in this assay (**Supplementary Figure S2B**). Conversely, fractionation on a gel filtration column showed reduced binding to Sgo1 and Sgo2 but again, does not allow us to distinguish between Sgo1- and Sgo2-specific functions of Set (**Supplementary Figure S2C**). We conclude that, at least i*n vitro*, Setβ binds Sgo1 and Sgo2 through interaction domains near both its N-terminus and C-terminus.

To address the *in vivo* importance of these interaction domains for Set function, oocytes were injected with mRNAs encoding GFP-Setβ ΔSgo N-term or GFP-Setβ ΔSgo C-term and induced to resume meiosis. Whole oocytes were fixed in metaphase I to assess chromosome alignment. Both Setβ mutant constructs localized to the centromere/kinetochore region, but neither rescued the chromosome alignment defects observed in *Set^-/-^* oocytes, unlike Setβ wild type (**Figure 2A and Figures 3C-E**). In mouse oocytes, Sgo2, rather than Sgo1, is required for error correction and thus, chromosome alignment^39, 40^. Together with our interaction data, we conclude that Set needs to interact with Sgo2 for proper chromosome alignment.

### Set function in chromosome alignment does not depend on PP2A-B56 binding

Set as an inhibitor of PP2A has been proposed to regulate Aurora B activity by inhibiting PP2A-B56^17, 21, 41, 42^. The Set C-terminus contains a conserved motif corresponding to the SLiM (short linear motif) for PP2A-B56 (**Supplementary Figure S3A**)^43–45^. Consistent with this, AlphaFold predicts that the SET SLiM binds to the B56 SLiM-binding pocket in a manner similar to that of another SLiM-containing protein, BUBR1, whose interaction with B56 has been well characterized (**Supplementary Figure S3B**)^43, 46, 47^. Indeed, *in vitro* assays with purified components showed that the PP2A-B56γ holo-complex, comprising all three subunits, interacted with Setβ, but not when the SLiM of Set was mutated (Setβ AxxAxA) (**Figure 3F, Supplementary Figure S3C**).

We investigated whether expression of Set carrying a mutation in the PP2A-B56 SLiM rescued the alignment phenotype of *Set^-/-^* oocytes. Contrary to expectations, Set’s direct interaction with PP2A-B56 was not required for chromosome alignment in meiosis I, as GFP-Setβ AxxAxA was still able to rescue chromosome alignment (**Figures 3E and 3G**). Thus, Set mediates this function in a Sgo2-dependent but PP2A-B56 SLiM-independent manner. Notably, the SLiM mutant also localized correctly to the centromere/kinetochore; therefore, failure to rescue alignment is not due to mislocalization of GFP-Setβ AxxAxA (**Figure 3D**). We speculate that the interaction between Set and PP2A-B56 regulates one of Set’s other multiple functions but is not necessary for chromosome alignment in meiosis.

### Bivalent segregation defects in *Set^-/-^* oocytes

Our study was initially motivated by the observation that Set plays a role in sister chromatid segregation in meiosis II, as reported by us and others^12, 23^. However, the transient knock-down approaches used in these previous studies did not allow precise determination of Set’s role in meiosis I, due to inefficient knock-down during the first division. The meiosis II phenotype-failure to correctly segregate sister chromatids in anaphase II-was weak and highly variable due to the transient approach being used^12^.

To assess how chromosome segregation in meiosis I is affected by Set loss, we analyzed chromosome spreads of control and *Set^-/-^* oocytes in metaphase II, i.e., after completion of meiosis I (**Figure 4A**). Importantly, unlike control oocytes, *Set^-/-^* oocytes still contained some bivalents (paired chromosomes) among the expected dyads (paired sister chromatids). Occasionally, bivalents appeared partially separated or held together along segments of chromosome arms (**Figures 4B and 4C**). The percentage of bivalents per oocyte was close to 20% (**Figure 4C**), and this phenotype was accompanied by sister chromatid pairs still connected along chromatid arms (“zipped dyads,” **Figure 4D**). Thus, without Set, oocytes incompletely separate chromosome arms in meiosis I. Consequently, the previously observed meiosis II phenotype (failure to segregate all dyads correctly in anaphase II) may have resulted from a general failure to remove Cohesin rather than a failure concerning solely pericentromeric Cohesin in meiosis II.

**Fig. 4.**
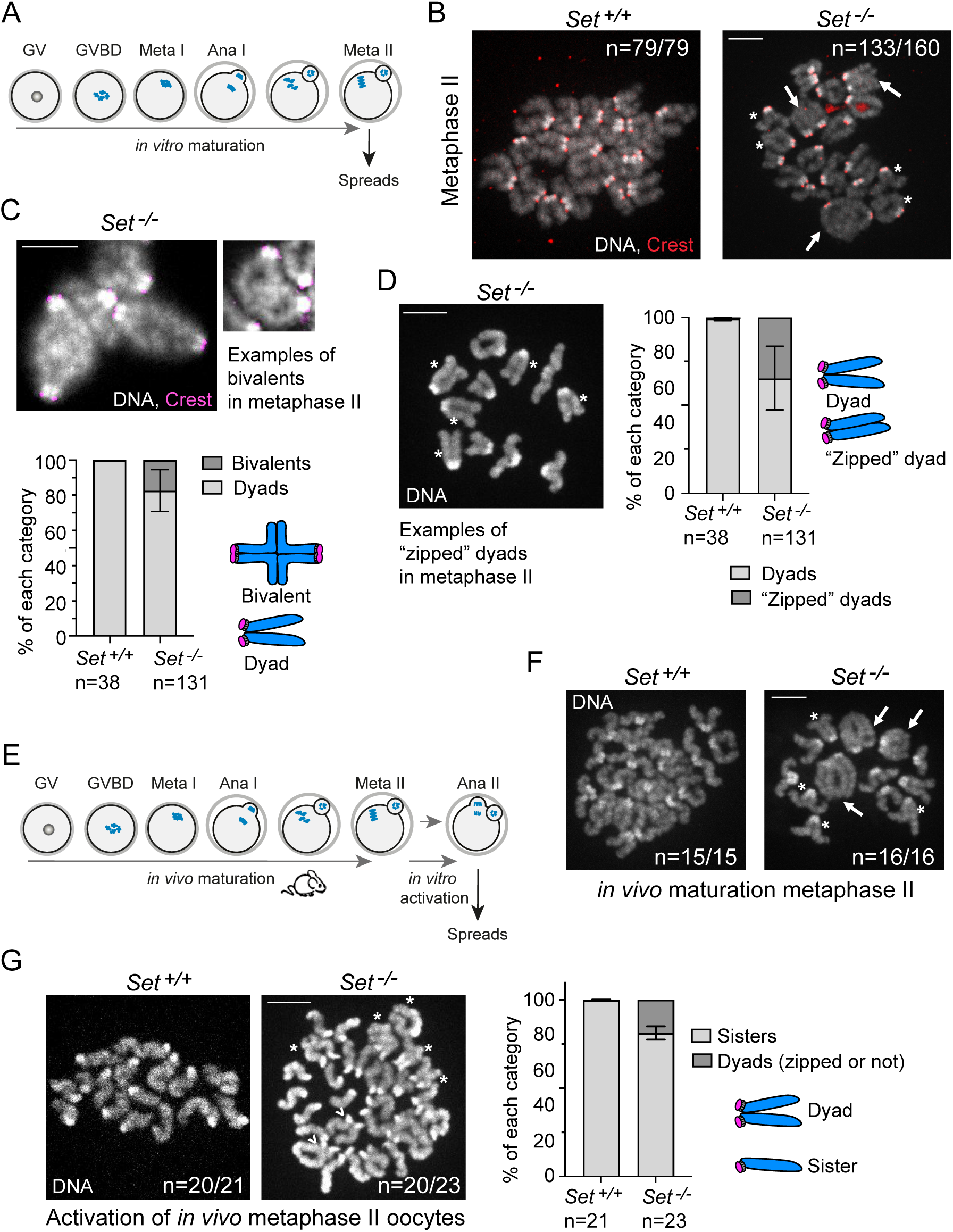
) Chromosome arms do not segregate efficiently in oocytes devoid of Set. **A** Scheme of experimental set-up of experiments in panel B-D. Oocytes were harvested at GV stage and matured *in vitro* until metaphase II. Metaphase II oocytes were used for the preparation of chromosome spreads. **B** Chromosome spreads in metaphase II *Set^+/+^* and *Set^-/-^* oocytes, stained with anti-Crest and Hoechst to mark centromeres/kinetochores (red) and DNA (grey), respectively. Arrow indicates a bivalent (chromosome pari that has not separated), asterisk indicates dyads (paired sister chromatids) still held together along chromosome arms. **C** Enlarged images of bivalents (arrows) of *Set^-/-^* metaphase II spreads obtained and stained as described in (B). For better visualization, chromosomes are shown in grey scale. Below, percentage of bivalents per oocyte. n indicates number of oocytes analyzed. **D** Enlarged images of dyads (asterisk) of spreads from *Set^-/-^* oocytes, stained with Hoechst. For better visualization, chromosomes are shown in grey scale. Below, percentage of “zipped” dyads per oocyte. n indicates number of oocytes analyzed. **E** Scheme of experimental set-up in panels F and G. Metaphase II oocytes were harvested from hormonally stimulated mice after *in vivo* maturation. Oocytes were activated *in vitro* through treatment with strontium, to induce anaphase II onset. **F** Chromosome spreads of *Set^+/+^* and *Set^-/-^* oocytes in metaphase II before activation, stained with DAPI to visualize chromosomes. The number of oocytes corresponding to the representative images shown from the total number of oocytes is indicated as n/total n. *Set^-/-^* oocytes either contained some bivalents and zipped dyads, or only zipped dyads. **G** Chromosome spreads of *Set^+/+^* and *Set^-/-^* oocytes after anaphase II, stained with DAPI to visualize chromosomes. Arrowheads specify dyads held together in the centromere region, and asterisk dyads held together along arms. On the right, quantification of dyads that did not separate correctly into sister chromatids. The number of oocytes corresponding to the representative images shown from the total number of oocytes is indicated as n/total n. All scale bars: 10 μm.

### Sister chromatid segregation failure in *Set^-/-^* oocytes

We examined whether *Set^-/-^* oocytes exhibited defects in meiosis II sister chromatid segregation. The conditional knock-out strategy allowed us to obtain metaphase II *Set^-/-^* oocytes that had matured *in vivo*, ensuring that segregation errors were not due to prolonged *in vitro* culture. Control and *Set^-/-^* metaphase II oocytes were harvested and chemically activated to mimic fertilization and anaphase II onset (**Figure 4E**). Indeed, we observed that before activation, *Set^-/-^* oocytes contained zipped dyads and bivalents as previously observed for *in vitro* matured oocytes (**Figure 4F**). Additionally, *Set^-/-^* oocytes often failed to segregate dyads correctly in anaphase II. Importantly, sister chromatids that did not segregate were frequently held together along chromatid arms in addition to the centromere region (**Figure 4G**). This indicates that the failure to segregate sister chromatids is due to inefficient removal of Cohesin from both chromosome arms and the centromere region.

### Rec8 is inefficiently removed from chromosome arms in *Set^-/-^* oocytes

The chromosome figures observed in chromosome spreads from *Set^-/-^* oocytes in meiosis I and II suggested that arm Cohesin was not efficiently removed. Indeed, Rec8 was frequently retained at unusual locations on bivalents of *Set^-/-^* oocytes, often along chromosome arms proximal to the centromere region (**Figure 5A**). This indicates that Set may interfere with Cohesin cleavage in meiosis I.

**Fig. 5.**
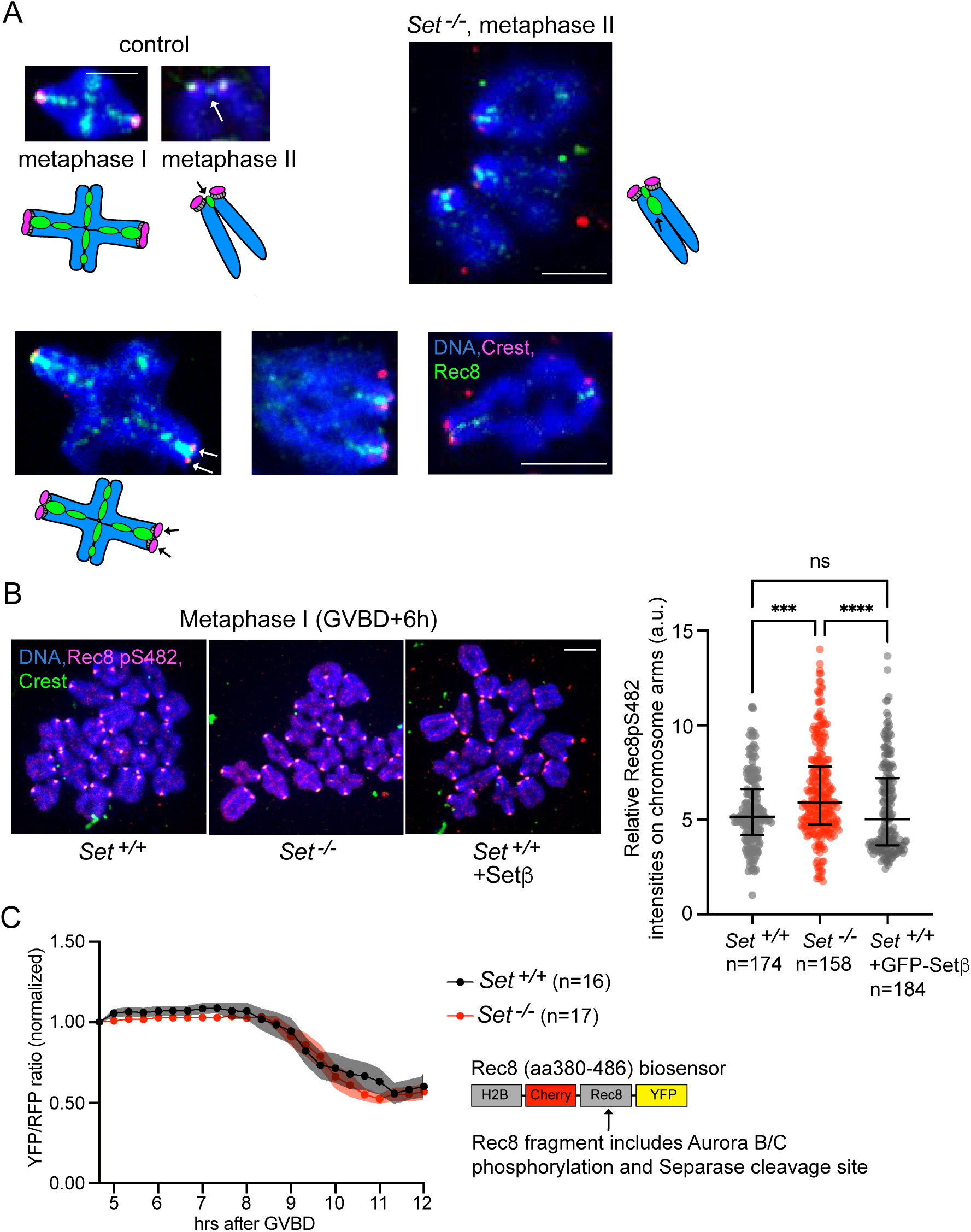
) Rec8 is retained on chromosome arms that do not segregate in Set-/- oocytes. **A** Examples of chromosomes in metaphase II spreads from *Set^+/+^* and *Set^-/-^* oocytes, stained with anti-Rec8 as well as anti-Crest and Hoechst to mark Rec8 (green), centromeres/kinetochores (red) and DNA (blue). In controls, examples of a metaphase I bivalent (chromosome pair) with Rec8 staining between sister chromatids of each chromosome (typical cross-shaped staining) and of a metaphase II dyad (sister chromatid pair) are shown. Pericentromeric Rec8 on a metaphase II dyad is indicated with an arrow. Several representative examples of dyads (right) and bivalents (bottom) in *Set^-/-^* metaphase II oocytes are shown. See schemes for explanation. On Set^-/-^ bivalents, white arrows (black arrows in the scheme) signify that separation of sister kinetchores has taken place. See text for details. **B** Chromosome spreads of metaphase I *Set^+/+^* and *Set^-/-^* oocytes (GVBD + 6 hours), stained with anti-Rec8 pS482 (red) and anti-Crest and Hoechst to mark centromeres/kinetochores (green) and DNA (blue), respectively. *Set^+/+^* oocytes were injected in GV stage with mRNA encoding GFP-Setβ, where indicated. On the right, quantification of the Rec8 pS482 signal normalized against background signal on chromosome arms. **C** *Set^+/+^* and *Set^-/-^* oocytes were injected with mRNA coding for a 380-486 aa fragment of Rec8 that is cleaved in a phosphorylation-dependent manner sandwiched between two fluorescent tags (YFP and Cherry) and localized to chromosomes due to Histone H2B (see scheme on the right). Rec8 cleavage is visualized due to change of colour of the sensor (from green to red), similar to the Separase activity sensor described in Fig 1F. Quantification of YFP/RFP ratio as a read-out of Separase activity towards Rec8. For quantifications in (B), n indicates the number of bivalents analyzed from the number of oocytes indicated on the left. Median and interquartile range are indicated, **** corresponds to p < 0,0001 and n.s. means not significant, using Kruskal-Wallis with Dunn’s multiple comparisons test. At least three independent biological repeats were analyzed for each condition. For quantifications in (C), each dot is mean and shaded areas indicate +/- SD. At least three independent biological repeats were analyzed for each condition. All scale bars: 10 μm.

Efficient cleavage of Rec8 on chromosome arms depends on Aurora B/C kinase-dependent phosphorylation of Rec8 at serine 482^48^. We hypothesized that Set might be required for efficient phosphorylation of Rec8 to facilitate cleavage. However, when comparing control and *Set^-/-^* oocytes in metaphase I, we detected higher, rather than lower, Rec8 phosphorylation with phospho-specific antibody staining (**Figure 5B**). Additionally, a Separase sensor containing a fragment of Rec8, including the Separase cleavage site and the phospho-site, was cleaved with comparable efficiency in control and *Set^-/-^* oocytes (**Figure 5C**)^48^. This is also in agreement with Aurora C still being localized to chromosome arms in *Set^-/-^* oocytes (**Figure 2C**). Thus, we conclude that oocytes lacking Set fail to remove all Rec8 from chromosome arms, but this is not due to inefficient Rec8 phosphorylation in the absence of Set.

### Set promotes arm Cohesin removal through N-terminal interaction with Sgo2

We used the same Set mutants as before to determine whether direct interaction with Sgo2 and/or PP2A-B56 is required for Set to promote arm Cohesin removal in meiosis I. Control and GV-stage *Set^-/-^* oocytes were injected with mRNAs encoding the different mutants, released into meiosis I, and fixed for chromosome spreads in metaphase II, as described above. Importantly, both GFP-Setβ and GFP-Setβ AxxAxA rescued bivalent segregation in meiosis I, demonstrating that arm Cohesin removal is promoted by Set independently of its direct interaction with PP2A-B56 through the identified SLiM (**Figure 6A**). GFP-Setβ ΔSgo C-term rescued bivalent segregation, whereas GFP-Setβ ΔSgo N-term did not (**Figure 6A**). This may be explained by *in vitro* gel filtration binding assays showing GFP-Setβ ΔSgo C-term to affect Sgo2 binding less penetrantly than GFP-Setβ ΔSgo N-term (**Supplementary Figure S2C**). Thus, contrary to chromosome alignment and cohesion protection, which require unhindered Set-Sgo2 binding, bivalent segregation may still proceed after partial loss of the Set-Sgo2 interaction.

**Fig. 6.**
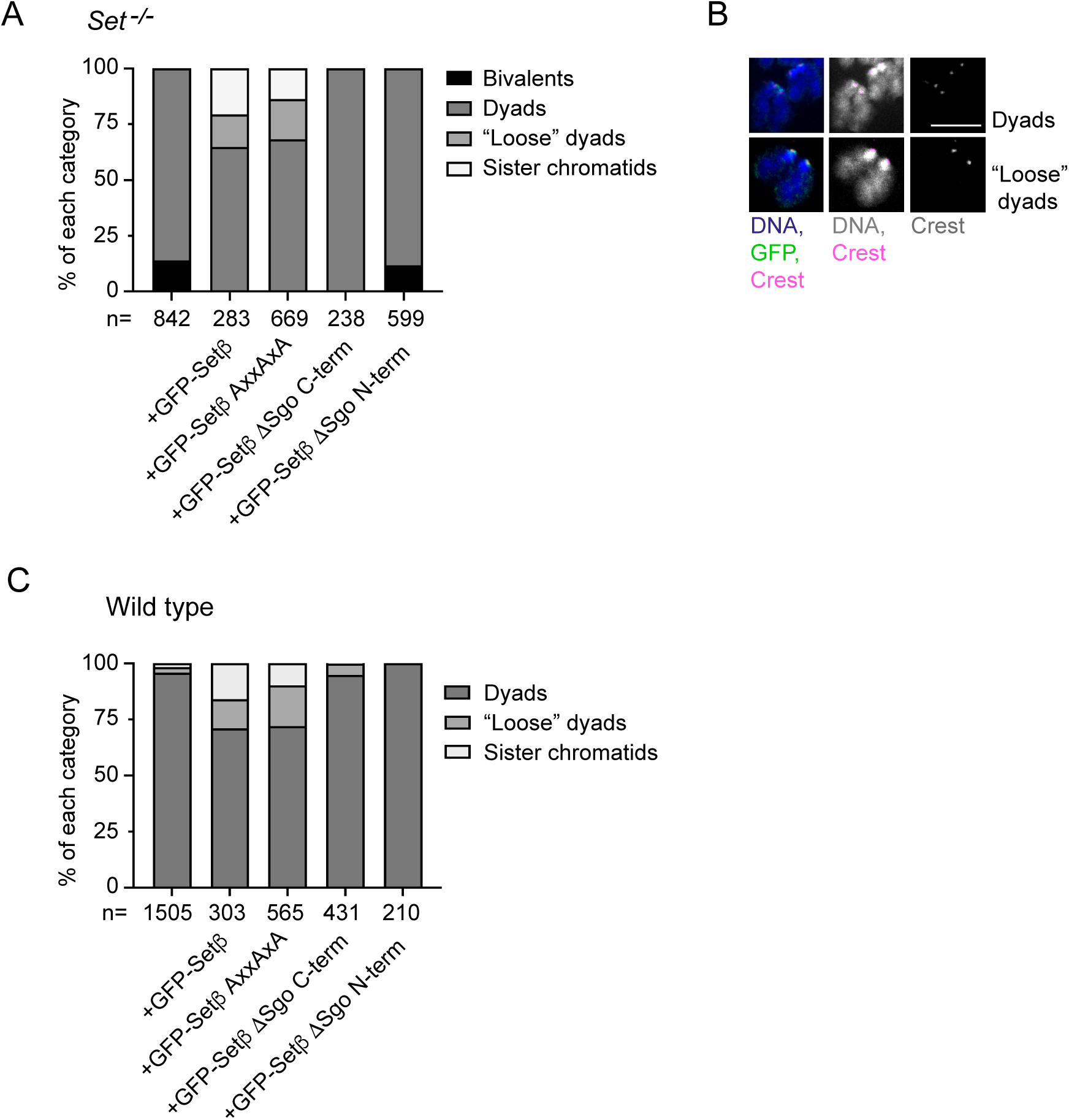
) Exogenous expression of Setβ mutants to assess requirements for bivalent segregation. Metaphase II spreads of wild type and *Set^-/-^* oocytes injected with mRNAs encoding the indicated GFP-tagged Setβ mutants. Spreads were stained with an anti-GFP antibody (exogenously expressed Set, green), anti-Crest (kinetochores/centromeres, red) and DAPI (DNA, blue). **A** Injection of *Set^-/-^* oocytes. **B** Phenotypes were classified as single sister chromatids, dyads, loose dyads and bivalents. Examples for dyads and loose dyads. **C** Injection of wild type oocytes. Of note, no bivalents were observed. n indicates the total number of chromosomes analyzed from at least three independent biological repeats per condition. Scale bar in (B): 10 μm.

### Setβ overexpression promotes precocious sister chromatid segregation through interaction with Sgo2 in meiosis I

Overexpression of Setβ in wild-type oocytes from the GV stage onward led to sparse instances of precocious sister chromatid segregation^23^ and of dyads that appeared separated but held in close proximity (“loose dyads”), likely due to incomplete decatenation by topoisomerase II^49–51^ (**Figures 6B and 6C**, **Supplementary Figure S1C**). To elucidate Set’s role in Cohesin removal around the centromere and on chromosome arms, we investigated whether interaction with Sgo2 and/or PP2A-B56 was required for Set to induce pericentromeric Cohesin removal and thus precocious sister chromatid segregation in meiosis I. Wild-type GV oocytes were injected with mRNAs encoding different Set mutants and induced to undergo meiosis I before being fixed for chromosome spreads in metaphase II. Expression of GFP-Setβ ΔSgo C-term and GFP-Setβ ΔSgo N-term did not induce precocious sister chromatid segregation or the presence of loose dyads, whereas GFP-Setβ AxxAxA promoted both (**Figure 6B**). Similar to wild-type oocytes, expression of GFP-Setβ and GFP-Setβ AxxAxA led to precocious sister chromatid segregation also in *Set^-/-^* oocytes **(Figure 6A)**. Since all mutants localized to the centromere region (**Figure 3D**), the failure of GFP-Setβ ΔSgo mutants to induce precocious sister chromatid segregation was not due to mislocalization. These results suggest that interaction with Sgo2 is necessary for Set to promote pericentromeric Cohesin removal, but unexpectedly, this occurs independently of Set’s interaction with PP2A-B56. Therefore, Set does not override Cohesin protection in meiosis I through direct interaction with PP2A-B56.

### Retention of phospho-Histone H1 but not Sgo1 on chromosome arms of meiosis I *Set^-/-^* oocytes

In mitosis, Set has been proposed to evict phospho-Histone H1 (H1p18, phosphorylated at serine 17 or 18, depending on the isoform) from chromosome arms, facilitating the removal of Sgo1 from arms and/or the centromere region for efficient Cohesin removal^16^. Sgo1 and H1p18 may thus be retained on chromosome arms in oocytes lacking Set. As shown above, interaction of Set with Sgo1 was much weaker than with Sgo2, and hardly perturbed by mutation of the N- and C-terminal Sgo interaction domains in Set, making it unlikely that Set promotes Cohesin cleavage by evicting Sgo1. However, knock-down of Sgo1 leads to some precocious sister chromatid segregation in meiosis I in mouse oocytes^52^. Therefore, we addressed whether H1p18, and consequently Sgo1, localize to chromosome arms and centromeres in meiosis I.

In wild type oocyte, endogenous H1p18 localizes exclusively to the centromere region in prometaphase I and metaphase I (**Figures 7A**). Importantly, in *Set^-/-^* oocytes, H1p18 accumulated on chromosome arms in addition to its centromeric localization (**Figure 7B**). Exogenous expression of GFP-Setβ in *Set^-/-^* oocytes reduced H1p18 levels on arms, indicating that Set removes H1p18 from chromosome arms (**Figure 7C**). To understand if increased phospho-H1 levels in *Set^-/-^* oocytes lead to retention of Sgo1, we evaluated Sgo1 localization on chromosome spreads in early prometaphase I (GVBD + 2 hours) and metaphase I (GVBD + 5–6 hours) in control and Set knock-out oocytes. No Sgo1 staining was detected on chromosome arms at any stage of meiosis I. Slightly lower Sgo1 staining was observed around centromeres/kinetochores in the absence of Set (**Supplementary Figure S4A and S4B**). The absence of Sgo1 from chromosome arms indicates that Sgo1 is most likely not evicted by Set in meiosis I.

**Fig. 7.**
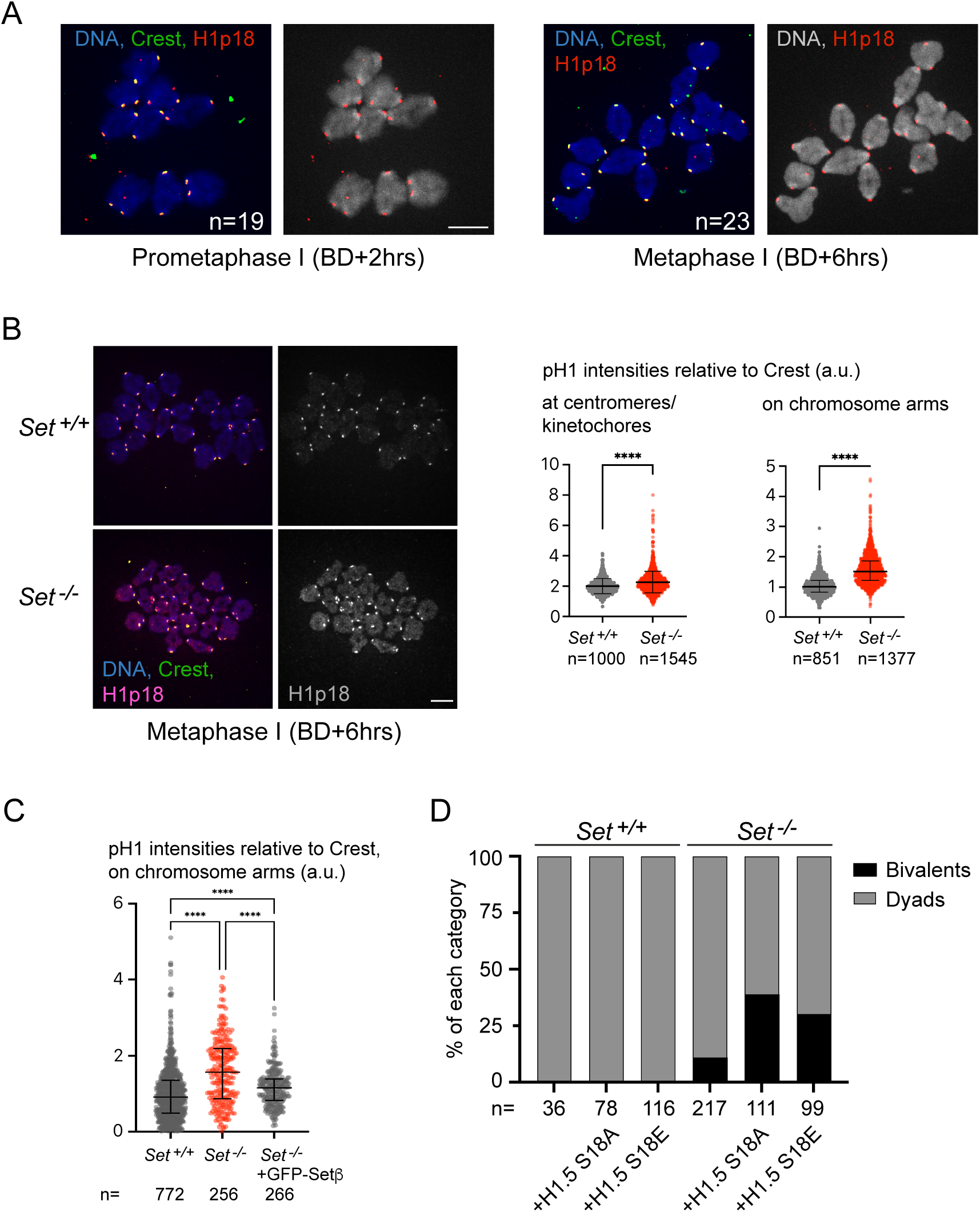
) Set is required to restrain Histone H1p18 localization to the centromere. **A** Localization of H1p18 on prometaphase I and metaphase I spreads from wild type oocytes. Spreads were stained with anti H1p18 antibody (red), anti-Crest antibody (centromeres/kinetochores, green) and Hoechst (DNA (blue or grey). Hours (hrs) after GVBD are indicated, n indicates the number of oocytes that were analysed for each timepoint. **B** Spreads were prepared and stained as described in (A), using metaphase I (GVBD+6 hrs) *Set^+/+^* and *Set^-/-^* oocytes. On the right, quantifications of the H1p18 signal relative to Crest, on centromeres/ kinetochores and on chromosome arms, respectively. n indicates the number of bivalents analysed. **C** Quantification of H1p18 levels on chromosome arms from metaphase I spreads of *Set^+/+^* and *Set^-/-^* oocytes. mRNA encoding GFP-tagged Setβ was injected were indicated. **D** Quantification of bivalents and dyads on chromosome spreads of metaphase I *Set^+/+^* and *Set^-/-^* oocytes, injected with either H1.5 S18A or H1.5 S18E. For quantifications in (C) and (D), n indicates the number of bivalents used for quantifications, from at least three independent biological repeats per condition. Median and interquartile range are indicated. **** corresponds to p < 0,0001, using Mann-Whitney U test in (C). **** corresponds to p < 0,0001 and ** to p = 0,0058, using Kruskal-Wallis with Dunn’s multiple comparisons test (D). For quantifications in (E), n indicates the number of oocytes analyzed, from at least three independent biological repeats per condition. All scale bars: 10 μm.

When we compared the sequence around the phosphorylation site in different histone H1 isoforms, we found that only Histone H1.3, H1.4, and H1.5 (in the mouse also called H1d, H1e, and H1b, respectively) contain a conserved sequence motif recognized by our anti-H1p18 antibody. To determine whether phosphorylation of H1 on Serine/Threonine 18 influences arm cohesion removal, we overexpressed phosphomimicking and non-phosphorylatable H1.5 from GV stage onward in wild-type and *Set^-/-^* oocytes and performed chromosome spreads in metaphase II. Whereas *Set^+/+^* oocytes showed no phenotype, *Set^-/-^* oocytes exhibited a more severe impairment in chromosome segregation upon expression of either GFP-H1.5 S18E or GFP-H1.5 S18A, than *Set^-/-^* oocytes alone (**Figure 7D**). The increased number of bivalents in metaphase II spreads indicates that oocytes become sensitive to elevated levels of both H1.5 S18E and H1.5 S18A only in the absence of Set, further demonstrating that Set counteracts H1.5 in oocyte meiosis. This aligns with the increased levels of endogenous H1p18 on chromosome arms in Set knock-out oocytes and suggests that Set is required to evict a distinct fraction of H1 from chromosome arms for efficient Cohesin removal by Separase. However, phosphorylation of H1.5 at serine 18 itself does not appear to be the key factor preventing proper chromosome segregation.

### Set is required to evict H1p18 and substitute it with the meiotic H1foo variant

We were puzzled by the observation that both non-phosphorylatable and phosphomimicking H1.5 increased the prevalence of bivalents in *Set^-/-^* oocytes. To explain this, we hypothesized that Set may specifically evict linker histones that are phosphorylated on this conserved phosphomotif. Consistent with this idea, this motif is absent in linker histones predominantly expressed in mouse oocytes, Histone H1.0 and H1foo (also called H1^0^ and H1oo, respectively). H1foo is an oocyte-specific linker histone and is presumably incorporated into chromatin in oocytes instead of somatic cells, where H1.1–H1.5, and H1.x prevail^53^. Set may therefore contribute to incorporation of H1foo into chromatin upon GVBD to replace somatic linker histones. We found that in wild type oocytes, H1foo is not associated with chromatin in GV oocytes before meiosis resumption (**Figure 8A, Supplementary Figure S5A**). Upon incubation of GV oocytes in culture medium, H1foo becomes even less concentrated in the GV than in the cytoplasm (**Supplementary Figure S5B**). H1p18 is found in the GV but not co-localizing with total H1 at chromocenters and DNA stained with DAPI (**Figure 8B**). Once oocytes resume meiosis I, H1p18 remains visible only around the centromere region (**Figure 7A**), but H1foo is now recruited to chromatin (**Figure 8C**). Indeed, when comparing prometaphase I *Set^+/+^* and *Set^-/-^* oocytes, we observed a significant reduction in chromatin-associated H1foo, despite identical cytosolic protein levels in GV oocytes (**Figures 8C and 8D**). Therefore, the decrease in chromatin-associated H1foo in the absence of Set is not due to lower H1foo protein levels. Set is thus important for establishing the correct chromatin environment after GVBD in oocyte meiosis, and its absence disrupts meiotic Cohesin cleavage.

**Fig. 8.**
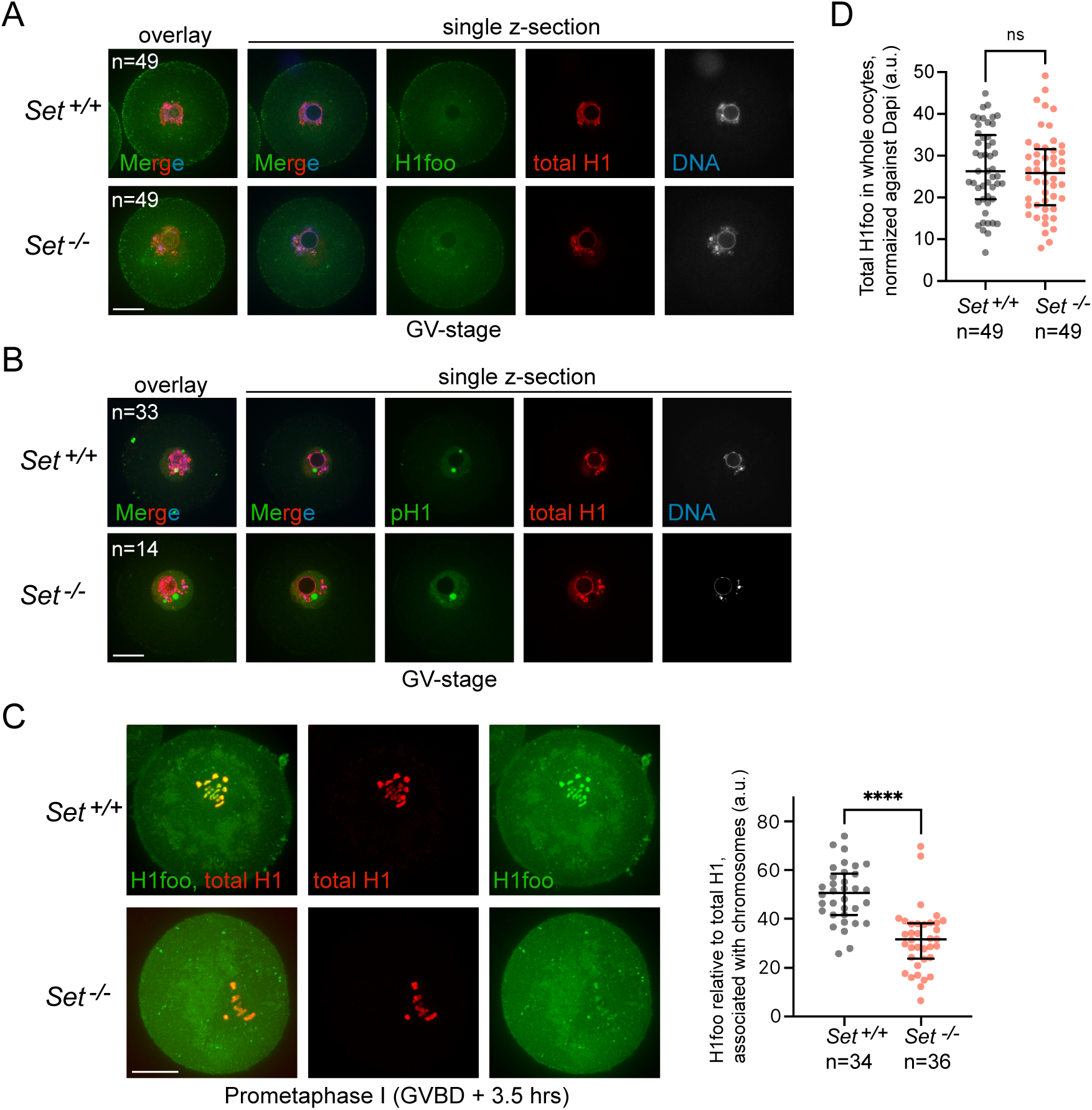
) In absence of Set, H1foo is not efficiently incorporated into chromatin after resumption of meiosis. **A** Whole mount *Set^+/+^* and *Set^-/-^* oocytes were fixed in GV stage and stained with anti-H1 antibody (red) and anti-H1foo antibody (green). Shown are the overlay of 20 z-sections of 2 μm of both channels (left) and a single z-section of each channel at the level of the GV with nucleolus (no staining). **B** Whole mount *Set^+/+^* and *Set^-/-^* oocytes were fixed in GV stage and stained with anti-H1 antibody (red), anti-H1p18 antibody (green), and DAPI (DNA, blue). Shown are the overlay of 20 z-sections of 2 μm of all three channels (left) and a single z-section of each channel at the level of the GV with nucleolus (no staining). **C** Whole mount *Set^+/+^* and *Set^-/-^* oocytes were fixed in prometaphase I at GVBD + 3.5 hours and stained with anti-H1 antibody (red) and anti-H1foo antibody (green). Shown are the overlay of 20 z-sections of 0.5 μm of both channels, and the overlays of the individual channels. **D** Quantification of total H1foo protein levels (cytoplasm and GV), normalized to DAPI. Median and interquartile range are indicated. **** corresponds to p < 0,0001, using Mann-Whitney U test in (B) and (D). n indicates the number of oocytes analyzed, from at least three independent biological repeats per condition. All scale bars: 20 μm.

## Discussion

Our work aimed to determine the roles of Set for mammalian oocyte meiosis, focusing on two key aspects of cell division: chromosome alignment and Cohesin cleavage. We generated a mouse model to obtain the oocyte-specific knock-out of Set, overcoming the limitations of transient knock-downs used in previous studies, which were poorly compatible with mutant rescue experiments focusing on specific molecular features of Set^12, 23^. We found that Set is essential for proper chromosome alignment in meiosis I. In its absence, misaligned chromosomes and missegregations are observed. We also discovered that Set promotes Cohesin cleavage by Separase, in both meiosis I and meiosis II. Importantly, the meiotic roles of Set do not require the PP2A-interaction domain that we have identified here, indicating that its direct interaction with PP2A is not required to promote proper chromosome alignment and Cohesin cleavage.

### Chromosome alignment

Aurora B/C-dependent substrate phosphorylation destabilizes erroneous, tensionless attachments, promoting the establishment of proper attachments. In mitosis, Set has been proposed to contribute to metaphase chromosome alignment through a PP2A inhibitory activity expected to facilitate Aurora B phosphorylation and activation^17, 21^. We show here that in oocytes, Set is also essential for chromosome alignment, and that error correction is weakened in the absence of Set. Phosphorylation of the Aurora B/C substrate Hec1 is reduced, coincidentally with a decreased Aurora C localization to centromeres that we observed (**Figure 2B and C**). Like in mitosis, Set has to interact with Sgo2 to rescue the chromosome alignment phenotype in Set knock-out oocytes^17^. However, a Set mutant lacking the PP2A-B56-binding SLiM can still rescue the chromosome alignment phenotype in Set knock-out oocytes. Thus, Set promotes alignment in a PP2A-B56-independent manner, at least in meiosis. Else, Set may inhibit PP2A through mechanisms other than direct binding. As Sgo2 is equally reduced at pericentromeres in Set knock-out oocytes, but loss of Sgo2 does not phenocopy loss of Set as far as chromosome alignment is concerned, we think that Sgo2 can both promote and inhibit Aurora B/C activity^39^. We conclude that Set promotes chromosome alignment in a Sgo2-dependent but PP2A-B56-independent manner in meiosis.

### Cohesin removal from chromosome arms

We found that without Set, Rec8 is not completely removed from chromosome arms in meiosis I (**Figure 6**). Absence of proper monopolar or bipolar spindle tension does not result in retention of Cohesin in meiosis I or meiosis II, respectively^54^, indicating that weakened error correction, resulting in less spindle tension, does not cause the cohesion defects observed in Set knock-out oocytes.

In mitosis, Set promotes Cohesin removal from chromosome arms through the prophase pathway^16^. In meiosis, in contrast, the prophase pathway does not remove Rec8 Cohesin once oocytes progress beyond prophase I arrest and only Separase-dependent cleavage of Rec8 is required for chromosome segregation^9^. Moreover, complete loss of Wapl, which is required for prophase-pathway dependent cohesin removal, has been shown to cause precocious sister chromatid segregation – a phenotype opposite to what would be expected for a mutation impairing Cohesin removal^4^. Therefore, the retention of Rec8 on chromosome arms in Set knock-out oocytes is unlikely to result from a defect of the prophase pathway.

In oocyte meiosis, we find that neither Sgo1 nor the Sgo2 pool required for Cohesin protection is increased on chromosomes upon loss of Set, contrary to our expectations based on mitotic data. Unlike Sgo2 which is found on chromosome arms in prometaphase I ^15, 55^, we never observed Sgo1 on chromosome arms (**Supplementary Figure S3**). The absence of Sgo1 as well as Sgo2 from chromosome arms in metaphase I suggests that eviction of either is not the mechanism by which Set promotes Cohesin cleavage.

Consistent with this, to promote Cohesin cleavage in meiosis I, Set does not require interaction with PP2A-B56, suggesting that chromosome segregation defects are unlikely to result from impaired inhibition of PP2A. However, Set still requires interaction with Sgo2. This raises a key question: How can Set promote arm Cohesin cleavage by Separase in metaphase I, if its binding and localization partner Sgo2 is absent from arms at this stage?

### Eviction of H1p18 allows incorporation of H1foo

Set can function as a histone chaperone, and indeed, Set was found to specifically evict H1pT18 from chromosome arms, creating favorable conditions for prophase pathway-dependent Cohesin removal^16^. Similar to mitosis, we observed accumulation of H1p18 on chromosome arms in metaphase I Set knock-out oocytes. Overexpression of phosphomimicking as well as non-phosphorylatable mutant H1.5 aggravated the phenotype of unsuccessful bivalent segregation in Set knock-out oocytes (**Figure 7**). This indicates that too much H1 (independently of the phosphorylation status of S/T18) interferes with Cohesin cleavage, but only in absence of Set. We think that in wild type oocytes enough endogenous Set is present to successfully evict excess H1.5, and that S/T18 phosphorylation is probably not the only modification required for eviction by Set.

If the prophase pathway is not required for chromosome segregation in oocyte meiosis, how can we explain that retention of H1p18 results in inefficient removal of Rec8-containing Cohesin from chromosome arms? We hypothesized that Set is necessary to create the optimal chromatin environment for Rec8 cleavage by Separase, and that in its absence, suboptimal conditions lead to inefficient Cohesin cleavage on chromosome arms. The H1p18 phosphorylation site is exclusive to somatic H1.3, H1.4, and H1.5, which are incorporated into chromatin until GVBD^56^. Conversely, the primary histones of oocytes undergoing the meiotic divisions are H1.0 and H1foo, which lack the sequence surrounding the phosphosite recognized by the H1p18 antibody^53^. Thus, staining with H1p18 antibody reveals the presence of somatic H1.3, H1.4, and H1.5, but not H1foo or H1.0.

When staining GV oocytes, we find that H1foo is not associated with chromatin. Phosphorylated - somatic-H1 is present within the GV but concentrated at locations distinct from the bulk of chromatin and chromocenters stained with DAPI, whereas a pan-H1 antibody stains all chromatin. Thus, we conclude that at GV, somatic H1s that are not phosphorylated on S/T18 are incorporated into chromatin, whereas H1foo is excluded. (We did not analyze H1.0 because it is not oocyte-specific and also present during male meiosis and the first embryonic mitosis.) Our results indicate that H1foo is incorporated only after oocytes resume meiosis I and undergo GVBD. Indeed, we observe strong recruitment of H1foo into chromatin as oocytes progress into prometaphase I. Notably, in the absence of Set, H1foo recruitment to chromatin is reduced, and as a consequence, H1p18 is aberrantly present on chromosome arms in metaphase I (**Figure 8**). We propose that Set, through interaction with Sgo2 (which is found on chromosome arms in prometaphase I^55^), promotes exchange of somatic H1s (phosphorylated on S/T18, but maybe also carrying other modifications) with H1foo on arms, once oocytes undergo GVBD.

### Importance of germ-line specific H1 variant for meiotic Cohesin cleavage

11 histone H1 variants exist in human and mouse, with 4 of them being specific for the germline, and H1foo being the only oocyte-specific variant. The somatic variants have distinct capacities to compact chromatin^57^ and cells show specific patterns of H1 variants associated with chromatin under various conditions (developmental state, differentiated or undifferentiated cells, during replication, depending on gene expression,…^57^). Probably because of redundancy between specific histone variants, knock-out mice for one specific variant do not show a phenotype^53^. The exact function of the oocyte specific H1 remains elusive, but it is attractive to speculate that in oocytes, H1foo (probably together with H1.0) generates a chromatin environment that favors Separase-dependent cleavage of Rec8. This model may explain why the Rec8 cleavage sensor, unlike endogenous Rec8, is cleaved with the same efficiency in presence or absence of Set (**Figure 8**). The sensor construct is localized to chromosomes with an H2B-tag, but its Rec8 moiety is not part of the Cohesin complex. We surmise that as part of the sensor, Rec8 is not exposed to the negative consequences of depleting H1foo, and is thus cleaved without delay.

### Cohesin cleavage in meiosis II

In meiosis II, Separase cleaves Cohesin at the pericentromere, which has been protected from cleavage in meiosis I. Initially, we thought that Set would promote solely pericentromeric Cohesin cleavage, by inhibiting PP2A at this location. However, here we found that Set knock-out oocytes contain bivalents and dyads with cohesive Cohesin being present also on the arms of some chromosomes. Upon anaphase II onset, sisters sometimes separate incompletely and are held together either along chromatid arms, in the centromere region, or both (**Figure 4**). The failure to separate arms in meiosis II is most likely due to inefficient Rec8 cleavage. We know from previous work that Separase has the capacity to cleave more than only pericentromeric Cohesin in meiosis II, so this phenotype is not due to the fact that there is too much Cohesin to cleave^15^. Thus, we think that arm separation in meiosis II fails for the same reasons than in meiosis I, namely, a suboptimal chromatin environment. The cause for the anaphase II phenotype is thus two-fold: incomplete arm separation in meiosis I that is carried over to meiosis II, and incomplete Cohesin cleavage in meiosis II. Thus, our data suggest that overall Cohesin cleavage is impaired by the absence of Set in both meiosis I and meiosis II. It remains to be determined if and how Set additionally promotes pericentromeric Cohesin cleavage. Precocious sister separation upon expression of Set mutants in meiosis I suggests that Set promotes pericentromeric Cohesin cleavage in a Sgo2-dependent manner, but again, independently of PP2A binding (**Figure 6**).

In conclusion, using an oocyte-specific Set knock-out and the temporal resolution of oocyte meiotic maturation, we provide critical insights into Set’s roles in chromosome alignment and segregation. It was unanticipated that the segregation phenotype was relatively mild, with only a few chromosomes or sister chromatids per oocyte failing to separate completely in meiosis I and II, respectively. However, it is important to note that histone chaperones often have overlapping and redundant functions, and we hypothesize that Set is not the sole chaperone preparing chromatin for Cohesin cleavage. Understanding Set’s role in oocyte chromosome and sister chromatid segregation is crucial for elucidating how aneuploidies arise, including in human oocytes. Interestingly, Set protein levels decrease with maternal age in oocytes^58^. Our work offers insights into how this may contribute to aneuploidies and the high error rates in human oocytes as women age.

## Acknowledgements

We thank Dr. Ingrid Vetter for providing support and her expertise in the generation of AlphaFold 3 predictions. We thank members of the Wassmann lab for discussions throughout the project. We are grateful to the administrative and informatics services at the Institut Jacques Monod and members of the animal house “Animalerie Buffon”. LK was the recipient of a PhD fellowship by FRM (Fondation pour la Recherche Médical ECO 20170637505), and a 4th-year prolongation fellowship (FDT 202001010942). SEJ received a 4th-year fellowship by FRM (FDT 202404018087), WEY a postdoctoral fellowship by the Association de la Recherche Contre le Cancer (ARC) and VS funding through an Erasmus fellowship. This work has received support from the investment program “France 2030” as part of the IdEx program (ANR-18-IDEX-0001) implemented by Université Paris Cité, under which the inIdEx project Formula is conducted. Work in the Wassmann lab was supported by the Fondation pour la Recherche Médical (Equipe FRM DEQ 20160334921 and Equipe FRM DEQ 202103012574), the Agence Nationale de la Recherche (ANR-19-CE13-0015 and ANR-23-CE13-0015-01) and furthermore by the French Government under the France 2030 program, managed by the National Research Agency (ANR), with reference ANR-24-PESF-0004. Work in the Musacchio lab is supported by the Max Planck Society, the European Research Council (ERC) Synergy Grant 951430 (BIOMECANET) and the DGF’s Collaborative Research Centre 1430 “Molecular Mechanisms of Cell State Transitions”.

## Material and Methods

### Mice

Mice were maintained under temperature, humidity, and light-controlled conditions under the authorization C75-05-13 at UMR7622 and authorization C75-13-17 at UMR 7592 in a conventional mouse facility, with food and water access *ad libitum*.and under 12h-light/12-hour dark cycle. The project was submitted to ethical review according to the French law for animal experimentation (authorization B-75-1308 and B-75-0513) in accordance with the application of the “3 R” rules and national guidelines, under license 5330. Adult CD-1 mice were purchased at 7 weeks of age (Janvier, France) and the *Set* conditional knockout line (C57BL/6 background) was bred in our animal facility to obtain *Setfl/fl Cre+* and *Setfl/fl* mice. Mice were PCR genotyped (primer sequences available upon request) and used between 9 and 16 weeks of age. For the *Set* conditional knockout line, litter mates (*Setfl/fl Cre-)* were used as controls. Except for hormonal stimulation (Figure 4F and 4G), mice were not involved in any procedures except for genotyping prior to being sacrificed by cervical dislocation between 8 and 16 weeks of age, to dissect ovaries and harvest GV stage oocytes. For *in vivo* maturation (Figure 4F and 4G) mice were stimulated by injecting 5 units of PMSG (Pregnant Mare Serum Gonadotropin, Abbexa LTD, abx260389), followed by 5 units of hCG (human Chorionic Gonadotropin, Abbexa LTD, abx260092) 48 hours later. 16-18 hours after hCG injection, metaphase II oocytes were harvested from oviducts.

### Mouse oocyte culture

Fully grown GV-stage oocytes were harvested from the ovaries of sexually mature female mice by puncturing with a needle. They were cultured in droplets of M2 medium (self-made or Merck Millipore, MR-015P) supplemented with 100 μg/ml dbcAmp (dibutyryl cyclic AMP; Sigma-Aldrich, D0260), which prevents resumption of meiosis I, covered with mineral oil (Sigma-Aldrich, M8410), and kept at 37°C. Follicular cells surrounding the oocytes were carefully removed by mouth pipetting with a narrow glass pipette. For further culture, the oocytes were washed and released in dbcAmp-free M2 medium. Only oocytes resuming meiosis I within 1h30 after release were kept and used for experiments. *In vivo* matured metaphase II oocytes were collected from stimulated mice by releasing cumulus-containing oocytes from the ampulla into M2 medium. Oocytes were dissociated in M2 medium with 0,125 mg/ml hyaluronidase (Sigma-Aldrich, H4272). Parthenogenic activation of *in vivo* cultured metaphase II oocytes was done by placing the oocytes into homemade M16 medium without CaCl_2_ for 30 minutes and then into the activation medium (M16 medium without CaCl_2_ supplied with 100 mM strontium chloride, Sigma-Aldrich 204463) for 30min, in a CO_2_ incubator. Then, oocytes were transfered into M16 medium and fixed for chromosome spreads 30 min to 1 hour later^59^.

### Jess capillary western blotting

Capillary-based western assay called the JESS Simple Western (ProteinSimple) was used to assess Set protein levels in extracts derived from 15 or 30 oocytes at GVBD + 4 hrs. Oocytes were lysed in 2 μL RIPA buffer, vortexed and snap frozen prior to the run. Graphical representation as pseudogel images was generated from peaks with Compass for SW software^27^.

### *In vitro* transcription and microinjections

Microinjection was done using self-made microinjection needles (using a magnetic puller, Narishige PN-31 and Next Generation Micropipette Puller P-1000, Sutter Instruments) connected to a FemtoJet microinjector pump (Eppendorf) with continuous flow, and a holding pipette (Eppendorf, VacuTip I EP51950000036-25, Vitrolife, 15331) and micromanipulator (Eppendorf), on a Nikon Eclipse Ti microscope. GV-stage oocytes were injected in dbcAmp supplemented medium with mRNA, transcribed using the mMessage mMachine T3 Kit (Invitrogen, AM1348) and purified using the RNase Mini Kit (Qiagen, 74104). Oocytes were kept arrested for 1-3 hours and checked for fluorescence before release. Separase sensor constructs to express Scc1 and Rec8 sensors have been published^28, 60^. Coding sequences for Setα, Setβ, Setβ mutants, H1.5 S18A and S18D have been PCR subcloned into a pRN3 plasmid with a GFP tag for *in vitro* transcription using the T3 promoter. Oocytes were kept arrested for 1-3 hours and checked for fluorescence before release.

### Oocyte fixation and immunofluorescence

Before fixation, oocytes were treated with acidic Tyrode’s solution (homemade) at 37°C for a few seconds to remove the *zona pellucida*. For chromosome spreads, oocytes were transferred into droplets of spreading solution (1% paraformaldehyde, Sigma-Aldrich 441244; 0.15% Triton-X100, Sigma-Aldrich T8787; and 3mM dithiothreitol, Sigma-Aldrich D9779, in distilled H2O) on microscope slides. The slides were allowed to dry at room temperature and kept at -20°C ^15^. For whole-mount fixation and spindle staining, oocytes were incubated 4-6 minutes in a cold treatment solution (80 mM PIPES, Euromedex 1124; 1 mM MgCl2, Euromedex 2189-C) at 4°C. For all whole-mount fixations oocytes were fixed for 30 minutes in BRB80 buffer (0.3% Triton-X100 and 1.9% formaldehyde, Sigma-Aldrich F1635) at room temperature, and permeabilized overnight in 1xPBS, 3% BSA, 0,1% Triton ^61^.

The following antibodies were used at the indicated concentrations: polyclonal rabbit Sgo2 antibody (gift from José Luis Barbero, 1/50), polyclonal rabbit anti Sgo1 antibody (A. Pendas, 1/150), rabbit polyclonal Histone H1 phospho-Thr17 antibody (CliniScience OAAF07576; 1/50), rabbit polyclonal anti Histone H1foo antibody (Abcam, ab71580, 1/50) mouse Histone H1 antibody (Santa-Cruz Biotech, sc-8030, 1/100) rabbit polyclonal anti-Rec8 (gift from Scott Keeney, 1/50), mouse monoclonal alpha-tubulin antibody (DM1A) coupled to FITC (Sigma-Aldrich, F2168; 1/100), rabbit polyclonal anti-Hec1 phospho-Ser55 (Euromedex GeneTex, GTX70017; 1/100), rabbit polyclonal anti-SET (Abcam, ab92872; 1/50), rabbit anti-Aurora C (Invitrogen, PA5-118973, 1/50), human Anti-Centromere Protein Antibody (ACA, Cliniscience 15-234-0001, 1/100) and human Crest auto-immune antibody (Cellon SA, HCT-0100; 1/100).

Secondary antibodies used were the following: donkey anti-rabbit CY3 (715-166-152, Jackson Immuno Research, 1:200), donkey anti-rabbit Alexa Fluor 488 (711-546-152, Jackson Immuno Research, 1:200), donkey anti-human Alexa Fluor 488 (709-546-149, Jackson Immuno Research, 1:200), donkey anti-mouse CY3 (715-166-151, **J**ackson Immuno Research, 1:200) and donkey anti-human CY3 (709-166-149, Jackson Immuno Research, 1:200). Hoechst 33342 (Invitrogen, H21492) was used during secondary antibody incubation at a concentration of 50 µg/ml to stain chromosomes, before mounting chambers in AF1 Citifluor mounting medium (Biovalley, AF1-100). Alternatively, slides were mounted in Vectashield mounting medium supplemented with DAPI (EUROBIO Scientific, H-1200).

### Image acquisition

Images of chromosome spreads were obtained using an inverted Zeiss Axiovert 200M or Nikon Eclipse Ti2-E spinning disk confocal microscope with a Plan-Apochromat 100x/1.4 NA (Zeiss) or Nikon 100x/1,45 Plan Apochromat oil immersion objective, coupled to an EMCCD camera (Evolve 512; Photometrics) combined with an MS-2000 automated stage (Applied Scientific Instrumentation), a Yokogawa CSU-X1 spinning disc, and a nanopositioner MCL Nano-Drive, controlled by Metamorph software. For each channel, 6 z-sections were recorded, separated by 0,4μm. Acquisitions of whole mount oocytes were done on the same microscopes using a a Plan-Apochromat 40x/1.4 NA oil immersion objective (Zeiss) or a Plan Fluor 40x/1.3 NA oil immersion objective (Nikon). For each GV oocyte, 20 z-sections were acquired, separated by 2 μm, for prometaphase I oocytes, 20 z-sections were acquired, separated by 0,5 μm. Live imaging was performed on the same Zeiss microscope using a Plan-Apochromat 40x/1.4 NA oil immersion objective, under temperature-controlled conditions and multistage acquisitioning piloted by Metamorph software. For each oocyte, 11 z-sections were recorded, separated by 3 μm, every 20 minutes for 10 hours. Whole oocytes in Figure 2A were imaged using an inverted Leica laser-scanning confocal microscope TCS SP5 II (Leica Microsystems, Heidelberg, Germany) equipped with a GaAsP hybrid detection system and a Leica 63x oil immersion objective (HCX Plan APO CS, NA 1.4). Scan speed was 400 Hz and z-section interval was 0.08 µm (Lam et al. 2016).

### Reverse transcription-quantitative PCR (RT-qPCR)

Reverse transcription was carried out with SuperScript III (Invitrogen) and random hexanucleotides for 1 h at 50°C on 1 μg RNA, quantified with a nanodrop (Thermo Scientific) from 20 GV-stage oocytes each. Real-time quantitative PCR (qPCR) was carried out on a Stratagene Mx3005p with Brilliant III SYBR Green master mix (Agilent) according to the manufacturer’s instructions. The primer sequences are available upon request.

### Preparation of expression vectors

A cDNA encoding human SETβ was generated using a cDNA fragment encoding SETα (codon optimized for *E. coli* expression and previously synthetized by GenScript). This fragment was subcloned in the pGEX6P-1 plasmid, and the segment encoding for the divergent N-terminal region of SETα was converted to SETβ by PCR and Gibson Assembly (New England Biolabs, Ipswich, US-MA). The final construct encodes a GST-(HRV3C site)-SETβ fusion protein and was validated by Sanger sequencing (Microsynth SeqLab, Göttingen, Germany). SETβ “AxxAxA” (L245A, I248A, E250A), ΔSgo N-term (D43K, E47K, E51K; adapted from mutations in^18^) and C-term (E206G, L217S;^17^) was generated by applying the QuikChange mutagenesis protocol. The cDNA encoding human PPP2R5C (splicing variant 3) was inserted in a pGEX6P-1 (codon optimized for *E. coli* expression and synthetized by GenScript). The final construct encodes a GST-(HRV3C site)-B56γ3 fusion protein. pGEX4T-1 expressing human PPP2R1A tagged with GST-(TEV site) was generated as in^62^. A sequence encoding His-PPP2CA was cloned in a pDEST8 vector for insect cells expression, with an uncleavable 8xHis-tag.

### Purification and assembly of PP2A-B56γ3 holoenzyme

#### Expression and purification of His-PPP2CA (PP2Ac)

PPP2CA was expressed and purified as previously described^63^. Briefly, 2 liters of Tnao38 insect cells were infected with baculovirus (at a 1:20 virus:cell ratio), cultured at 19°C for 4 days ^64^. Cells were lysed by sonication in lysis buffer (Hepes 50 mM pH 7.5, 150 mM NaCl, 5% vol/vol Glycerol, 2 mM TCEP) supplemented with DNAse (Roche) and Protease inhibitors (Serva Electrophoresis GmBH) and cleared by centrifugation for 45 min at 85000X*g* at 4°C. The resulting supernatant was collected, filtered with a 0.8 μm filter and loaded 2 x 5 mL Talon column (Cytiva). After extensive washes, the protein was eluted in 2 mL fractions with 250 mM Imidazole in lysis buffer. Fractions containing PP2Ac were selected using Bradford reagent (Thermo Fisher Scientific) were diluted with ion exchange buffer (50 mM Hepes pH 7.5, 50 mM NaCl, 1 mM TCEP, 5% vol/vol glycerol) and loaded on a 6 mL ResourceQ column (Cytiva). The column was washed until no absorbance at 280 nm was detected by the ÄKTA system (Cytiva) used. Protein was eluted by applying a gradient of salt up to 400 mM NaCl in ion exchange buffer. Fractions were collected, concentrated and loaded on a S75 10/300 size-exclusion chromatography column (Cytiva) equilibrated with lysis buffer. Fractions showing the highest protein enrichment were collected, pooled, flash-frozen in liquid nitrogen, and stored at -80°C.

#### Expression and purification of GST-PPP2R5C (B56) and GST-PPP2R1A (PP2Aa)

GST-PPP2R5C (B56) and GST-PPP2R1A (PP2Aa) were expressed and purified as previously described, with few modifications^63^. The constructs were individually expressed in BL21 Codon Plus (DE3)-RIL *Escherichia coli* (Agilent Technologies, #230240) respectively growing in Lysogeny Broth (PPP2R5C) or Terrific Broth (PPP2R1A). Expression was induced by adding 0.1 mM IPTG to a 1-Liter culture when OD was equal to 0.6-0.7 or 0.8-1.0, respectively. Cells were grown for 18 hours at 18°C and were resuspended in purification buffer (50 mM Hepes 7.5, 300 mM NaCl, 5% glycerol, 1 mM TCEP; for PPP2R1A NaCl was lowered to 250 mM) supplemented with DNAse (Roche) and Protease inhibitors cocktail mix (Serva Electrophoresis GmBH). All purification steps were performed on ice or at 4-8°C. Cells were ruptured by sonication and the lysate cleared by centrifugation for 30 min at 85000X*g*. The cleared lysate was loaded on 5 mL of pre-equilibrated (purification buffer) slurry of glutathione beads (Serva) and incubated under rotation for 2 hours. The flow-through was discarded by means of a gravity-flow column (Thermo Fisher Scientific, Waltham, US-MA) and the beads were washed with purification buffer. GST-PPP2R1A was eluted by a single step addition of 20 mM Glutathione in purification buffer. While bound to beads, GST-PPP2R5C was incubated overnight at 8°C with 1 mg of GST-PreScission protease (produced in house) to cleave the N-terminal GST tag and release the protein in purification buffer. The protein was then concentrated and injected onto a Superdex 200 16/600 size-exclusion chromatography column (Cytiva) pre-equilibrated in purification buffer. Fractions showing the highest protein enrichment were collected, pooled, flash-frozen in liquid nitrogen, and stored at -80°C.

#### Assembly of the PP2A holocomplex

No pre-assembly of the PP2A holocomplex was performed. Individual PPP2R1A, PPP2CA and PPP2R5C subunits at concentrations indicated in the conditions for the pull-down protocol were mixed with or without SETβ and incubated for 1 hour.

### Expression and purification of wild type SETβ, SETβ AxxAxA, N-term ΔSgo, and C-term ΔSgo mutants

SETβ protein was expressed in 1 L of BL21 Codon Plus (DE3)-RIL cells growing in Terrific Broth (TB). Expression was induced by adding 0.1 mM IPTG to the media and allowed to continue for 18 hours at 18°C. Cells were pelleted via centrifugation and stored at -80°C until protein purification. The pellet was resuspended in purification buffer (50 mM Hepes 7.5, 250 mM NaCl, 5% Glycerol, 1 mM TCEP) supplemented with DNAse (Roche) and protease inhibitor mix (Serva Electrophoresis GmBH). Cells were lysed by sonication and a cleared supernatant was obtained by centrifugation for 30’ at 85000X*g*. The cleared lysate was loaded onto 5 mL of pre-equilibrated (purification buffer) glutathione bead slurry (Serva Electrophoresis GmBH) and incubated under rotation for 2 hours. The flow-through was discarded by means of a gravity-flow column (Thermo Fisher Scientific) and the beads were washed with purification buffer. While bound to beads, GST-SETβ was incubated overnight at 8°C with 1 mg of GST-PreScission protease (produced in house) to cleave the N-terminal GST tag and release the protein in purification buffer. The resulting protein was then concentrated and injected onto a Superdex 200 16/600 size-exclusion chromatography column (Cytiva) pre-equilibrated in purification buffer. Fractions showing the highest protein enrichment were collected, pooled, flash-frozen in liquid nitrogen, and stored at -80°C.

### Pull-down assay

Proteins were diluted to 5 μM concentration using binding buffer (50 mM HEPES pH 7.5, 5% glycerol, 1 mM TCEP, and 0.05% Tween) to a total volume of 50 μL (for experiment in supplementary Figures 3B and S2B, protein concentration was lowered to 3 μM and 5 μM, and the binding buffer was used with increased NaCl concentration equal to 250 mM and 150 mM, respectively). After centrifugation at 16900X*g* for 30’, samples were mixed with 10 μL glutathione beads (Serva Electrophoresis GmBH) and incubated on ice for 1 h. At the end of incubation, 10 μL of the mix were removed and mixed with 40 μL of 2x SDS-PAGE sample loading buffer (input sample). To separate the beads from the unbound proteins, the samples were centrifuged, at 1000X*g* for 1 min at 4°C. The supernatant was removed, and the beads were washed two times with 500 μL binding buffer. After the last washing step, 50 μL of 2× SDS-PAGE sample loading buffer were added to the dry beads. The samples were analyzed by SDS-PAGE and Coomassie staining or by Western blotting. The latter was carried out using the TransBlot Turbo transfer system (BioRad Laboratories). A pan-SET antibody (Abcam; #ab181990, EPR12973) was used at 1:1000.

### Gelfiltration

Analytical SEC was performed on a Superdex 200 5/150 Increase (Cytiva), mounted on an ÄKTA™ micro system (Cytiva). In total, 50 µL samples at 3 µM single protein concentration have been mixed then centrifuged at 16,900× g for 15 min. After sample injection, the protein absorbance was monitored at 280 nm, and the proteins were eluted under isocratic condition, at a flow of 0.15 mL/min, in 100 µl fractions at 4 °C. Fractions were analyzed by SDS–PAGE and Coomassie Blue staining. Fractions from 1 mL elution volume to 2.3 mL have been loaded on SDS–PAGE gels, subsequently stained with Coomassie brilliant blue (produced in-house).

### Quantification and statistical analysis

ImageJ software (NIH) was used for all image analysis. On chromosome spreads, the mean fluorescence intensity of each channel was quantified on sum-projected images by drawing an 8-pixel circle around centromeres marked by Crest; the background signal for each point was subtracted. For Supplementary Figure S4, 13x13 pixels were drawn around centromeres and background-subtracted. On whole oocyte immunofluorescence, quantifications were done on one z-section at the level of the germinal vesicle and the nucleolus. 300-pixel squares were drawn including the GV and surrounding cytoplasm for quantification and background subtraction. In Figure 2C, quantifications were done with ImageJ 1.54p, using the TrackMate plugin^65^ to draw circles of 4 pixels in diametre of the ACA signal to quantify the Aurora C signal at that location, and background was substracted. For all quantifications, raw values were normalized to the control values of the same experiment to allow comparison between different experiments. All data plots and statistical analysis were done with GraphPad Prism 8.0 and 11.0 software. Quantification results were compared using nonparametric Mann–Whitney U-test when comparing two groups, and Kruskal-Wallis test with Dunn’s multiple comparison test when comparing more than two groups. For RT-qPCR, unpaired student t-test was used. For quantification of the pull-down assays, raw images were uploaded on ImageJ, and background was subtracted with default settings via the built-in option. ROIs of the same area were generated and the integrated density was used as signal intensity value. Set binding specific signal was calculated by first subtracting the unspecific signal coming from the negative control lanes, then by normalizing against the signal intensity of the bait. Set signal is provided as a fraction relative to the control condition (equal to 1.0).

### AlphaFold-based modeling and SLiM alignment

Alpha-Fold predictions of SET:B56 and BUBR1:B56 complexes shown in Supplementary Figure S3B were performed using AlphaFold version 3^66^ and displayed using Chimera UCSF^67^. Uniprot codes of the proteins used in the predictions: B56 (Q13362-1), SET (Q01105-2), BUBR1 (O60566-1). PP2A-B56 subunit colored in silver (BUBR1 model)/ light blue (SET model, SET SLiM peptide in coral red and BUBR1 SLiM peptide in green. Alignment in Supplementary Figure S3A was done using Jalview and Mafft with default settings as alignment tool^68^. Set sequences Uniprot codes: Q01105-2, Q9EQU5-2, F2Z4L4, Q28FE9, Q7ZUY0, P53997.

## Supplementary Figures

**Supplementary Figure 1.**
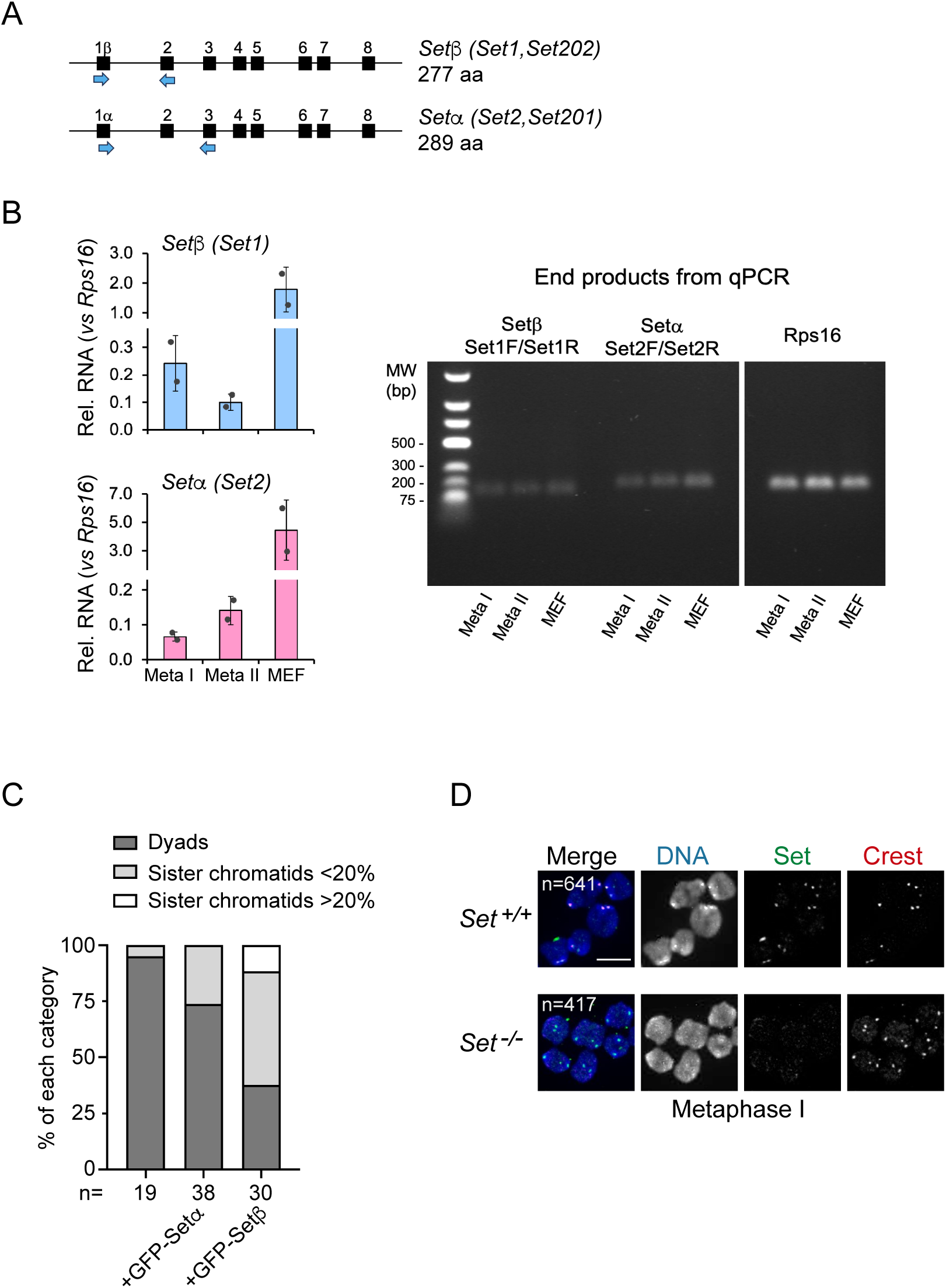
Both Setα and Setβ are transcribed in mouse oocytes, but Setβ induces a stronger phenotype when overexpressed. **A** Scheme of primer design to detect Setβ and α, respectively. **B** RT-qPCR using *Set^+/+^* and *Set^-/-^* oocytes in GV stage as well as Mouse Embryonic Fibroblasts (MEF), with primer pairs indicated in (A). PCR end-products were analyzed by gel electrophoresis. Rsp16 was used as internal control. **C** mRNAs coding for GFP-Set α or β were injected in wild type GV oocytes. Meiotic resumption was induced, and oocytes were fixed for chromosome spreads in metaphase II and stained with Crest and DAPI. The percentage of dyads and single sister chromatids was determined per oocyte. **D** Metaphase I chromosome spreads from *Set^+/+^* and *Set^-/-^* oocytes stained with anti-Set antibody (green), anti-Crest antibody to reveal kinetochores/centromeres (red) and Hoechst to stain DNA (blue). B) dots indicate individual experimental repeats, and unpaired student t-test was used. C) n indicates the number of oocytes analyzed from three independent biological repeats per condition, D) indicates the number of chromosomes analyzed from three independent biological repeats per condition. Scale bar: 10 μm.

**Supplementary Figure 2.**
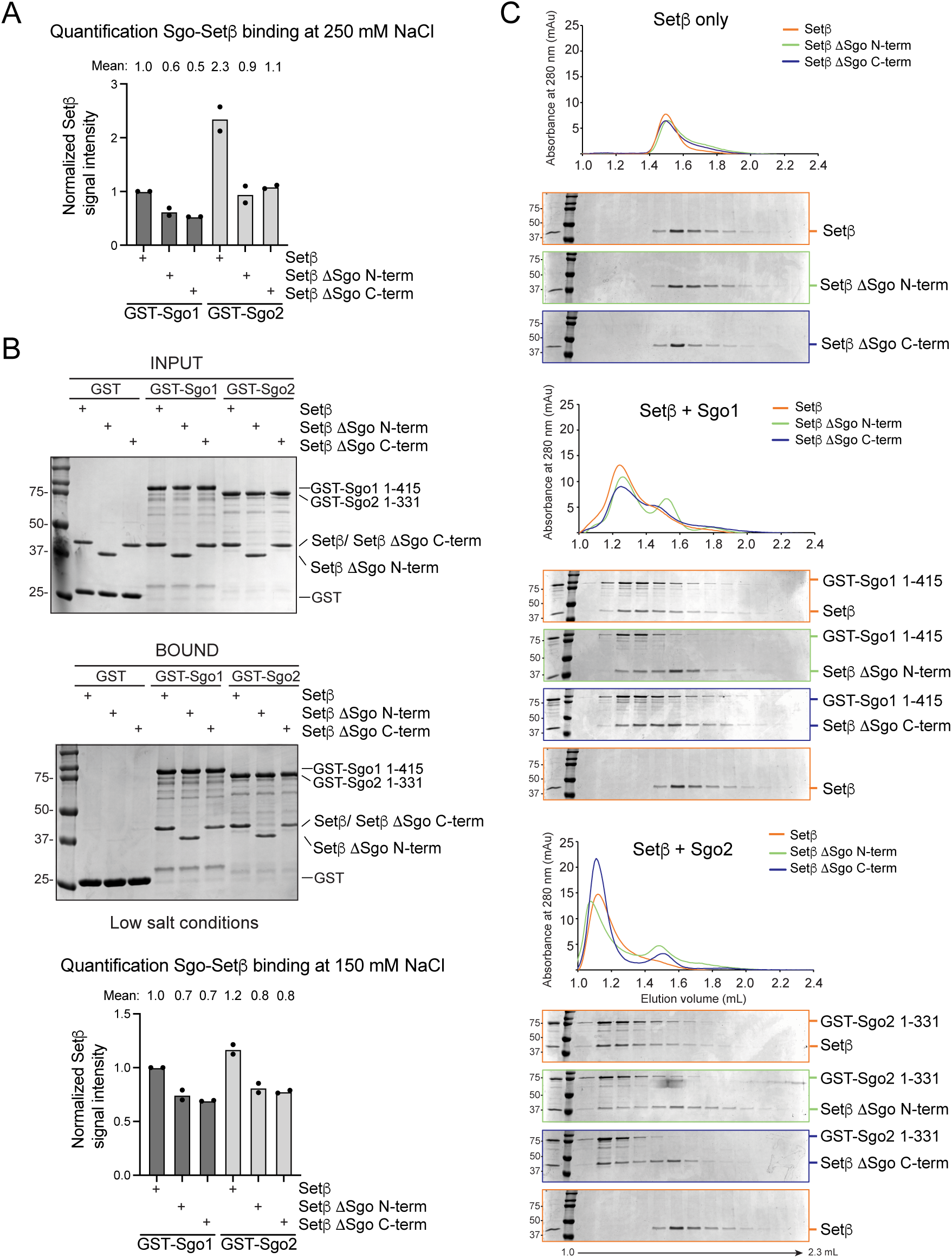
Mutants for N and C-terminal domains of Setβ do not allow to distinguish binding to Sgo1 and Sgo2. **A** Quantification of two repeats of pulldowns in (Figure 2B). **B** Binding assay under low salt conditions on Glutathione beads with GST-Sgo1 1-415, GST-Sgo2 1-133 (bait) and GST (negative control). Setβ wild-type, Setβ ΔSgo C-term or Setβ ΔSgo N-term were added in solution such as indicated. Beads were recovered by centrifugation, washed and analyzed by SDS-PAGE and coomassie blue staining. Below: Quantification of two repeats. **C** Setβ wild-type, Setβ ΔSgo C-term or Setβ ΔSgo N-term, GST-Sgo1 1-415 and GST-Sgo2 1-331 were diluted to 3 μM in 50 μL. Samples were centrifuged for 15 minutes at 16900g and then injected in a Superdex 200 5/150. Eluted fractions from 1 mL to 2.3 mL were run on an SDS-PAGE gel and stained with Coomassie Brilliant Blue. The experiment consists of two biological replicates. The Set signal correpsonds to the fraction relative to the control condition. For quantifications, dots represent repeats, means are indicated.

**Supplementary Figure 3.**
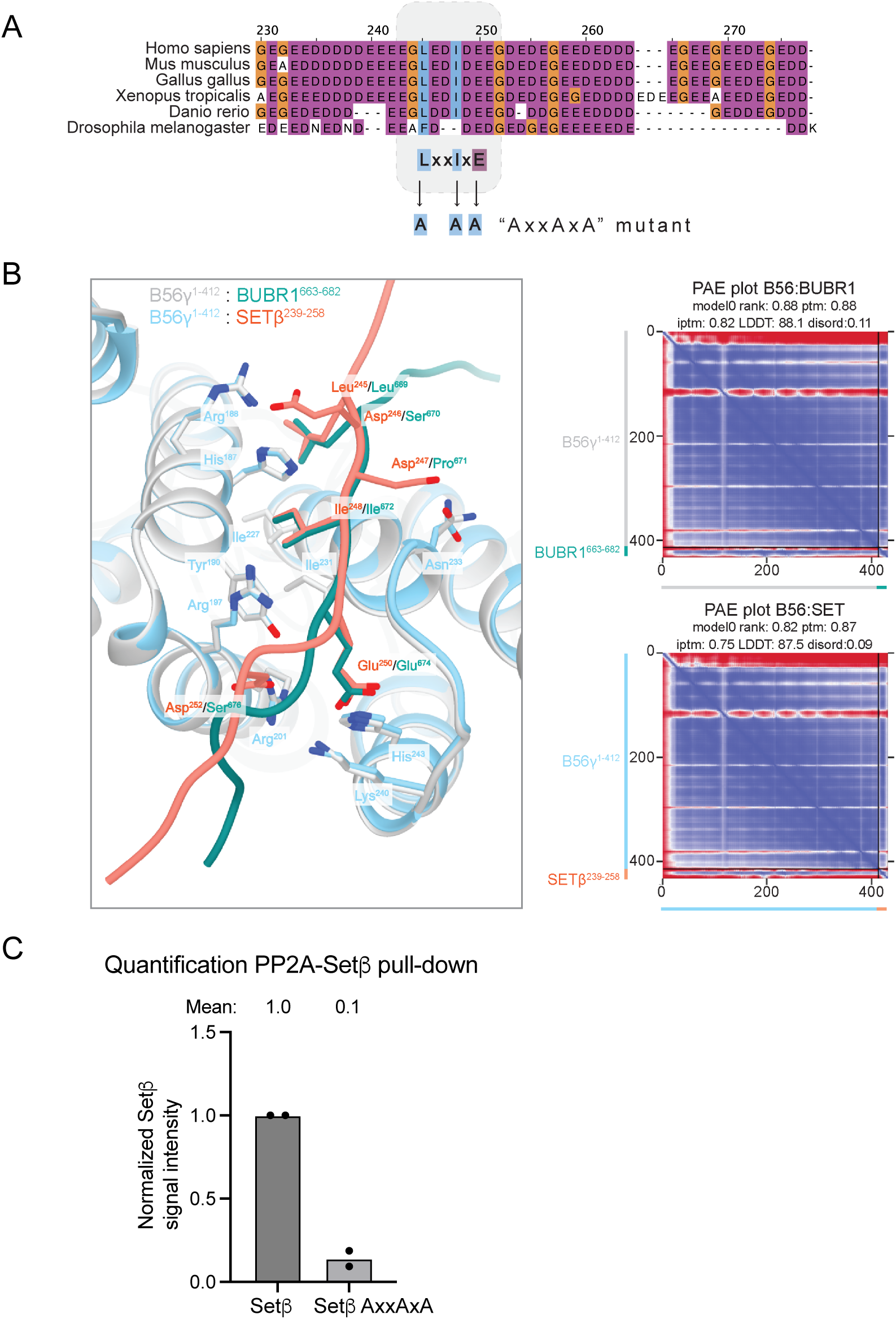
AlphaFold3 model of BUBR1 and Set SLiM interacting with binding pocket of PP2A-B56. **A** Conservation of PP2A-B56 SLiM in Setβ. (colored with the Clustal X style, where magenta indicates negatively charged residues, blue hydrophobic residues, orange glycines, white unconserved residues) **B** On the left: Superposed AlphaFold3 (AF3) ribbon models showing the SLiM binding pocket of B56 bound to BUBR1 SLiM (dark cyan) or Set SLiM (coral red). On the right: Predicted Aligned Error plots for the AF3 models shown on the left. **C** Quantification of two repeats of pulldown in (Figure 3F). The Set signal corresponds to the fraction relative to the control condition. For quantifications, dots represent repeats, means are indicated.

**Supplementary Figure 4.**
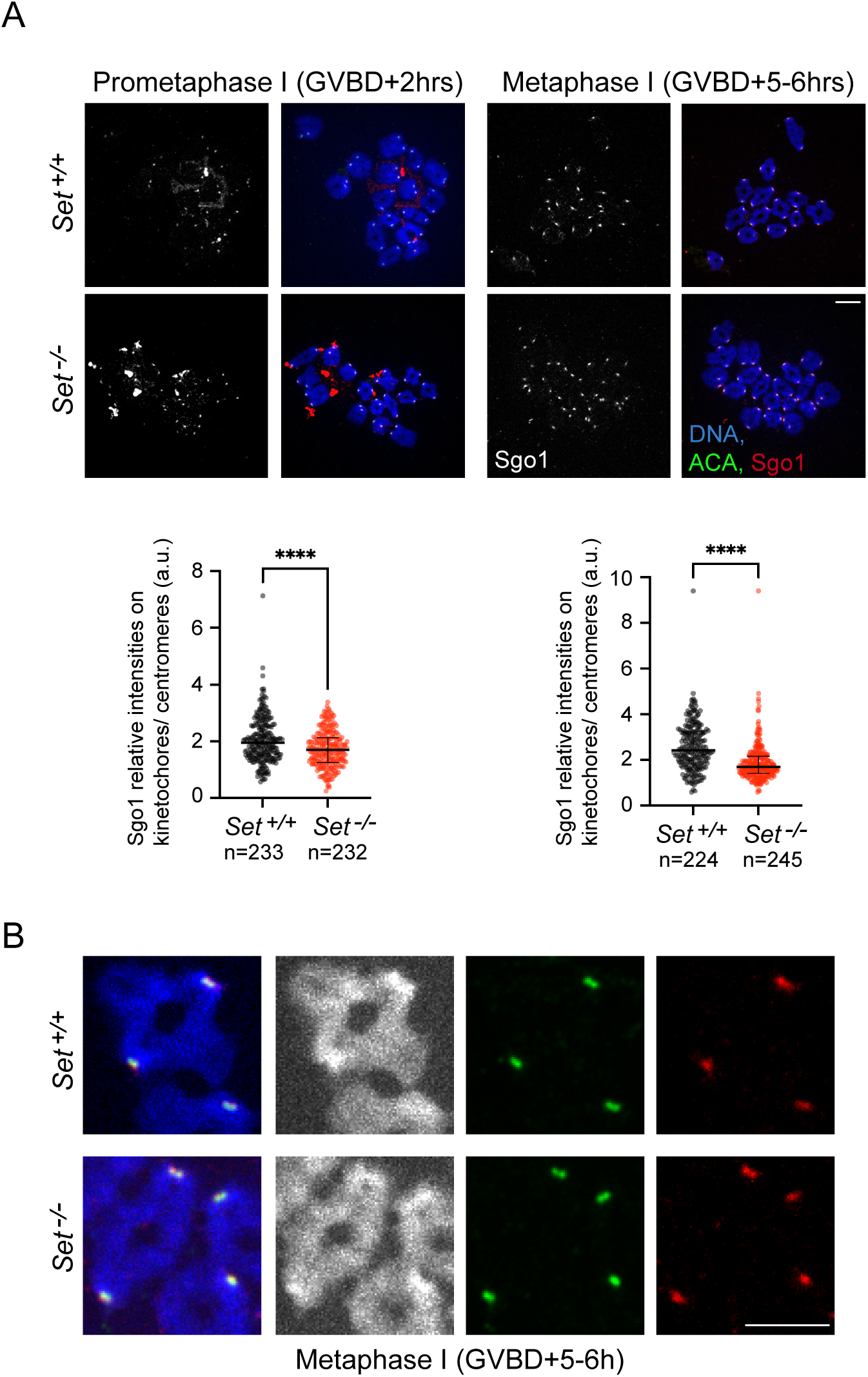
Sgo1 localization is not retained on chromosomes upon loss of Set. **A** Chromosome spreads of *Set^+/+^* and *Set^-/-^* oocytes in prometaphase I (GVBD + 2 hours) and metaphase I (GVBD + 6 hours), stained with anti-Sgo1 antibody (red), anti-ACA antibody to reveal kinetochores/centromeres (green) and DAPI to stain DNA (blue). Below, quantifications of the Sgo1 signal relative to ACA signal. n indicates number of kinetochore pairs analysed. **B** Zoom on selected bivalents from experiment in (A). For quantifications, n indicates the number of dyads analyzed, median and interquartile range are indicated and **** corresponds to p < 0,0001, using Mann-Whitney U test. At least three independent biological repeats were analyzed for each condition. All scale bars: 10 μm.

**Supplementary Figure 5.**
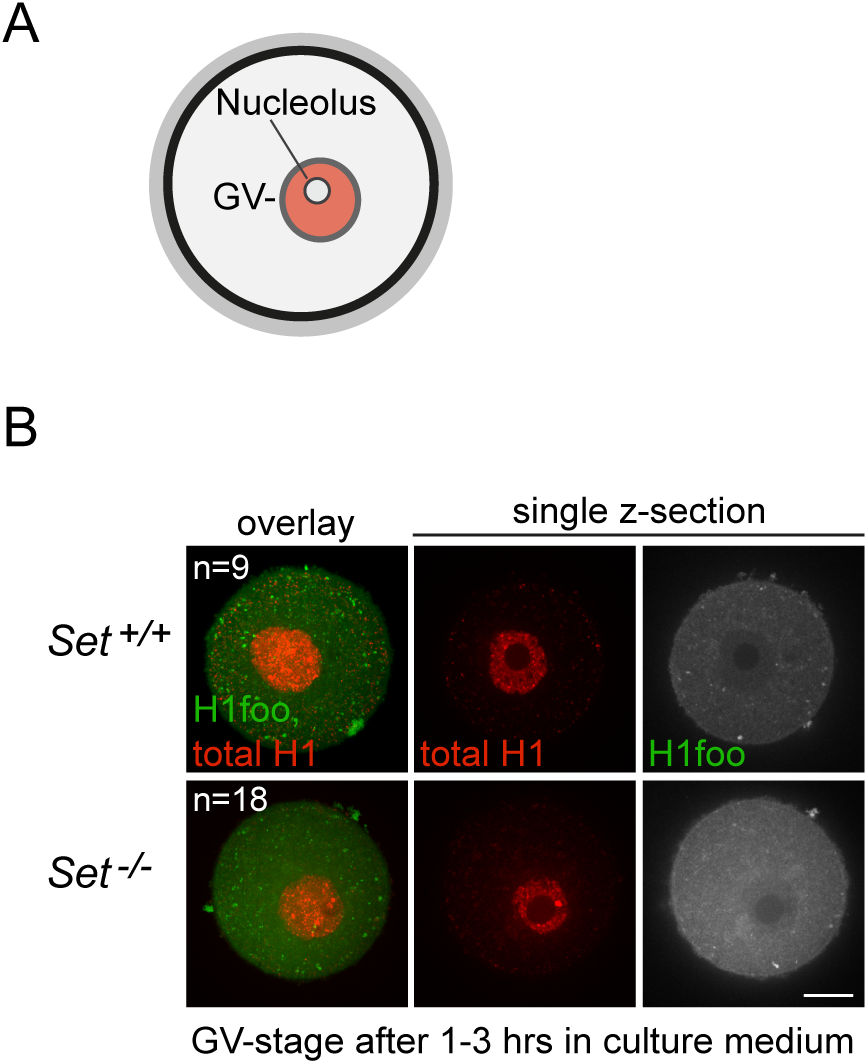
H1foo is not retained in the GV before resumption of meiosis I. **A** Scheme of a GV oocyte. The nucleolus inside the GV, which is visible on whole mount oocyte staining at GV stage in Figure 8, is indicated. **B** Whole mount *Set^+/+^* and *Set^-/-^* oocytes were fixed in GV stage after incubation in culture medium for 1-3 hours and stained with anti-H1 antibody (red) and anti-H1foo antibody (green). Shown are the overlay of 20 z-sections of 2 μm of both channels (left) and a single z-section of each channel at the level of the GV with nucleolus (no staining). n indicates the number of chromosomes analyzed, from three independent biological repeats per condition. Scale bar: 20μm.

